# Glucocorticoid rhythm disruption drives hyperinsulinaemia in mice through beta cell glucocorticoid receptor signalling

**DOI:** 10.64898/2026.06.03.730005

**Authors:** Jijo Wilson, Anna S. Arzeno, Sanjeev Sharma, Agnieszka Agas, Jack Lungstrum, Mary N. Teruel

## Abstract

**Aims/hypothesis:** Hyperinsulinaemia is typically viewed as a secondary, compensatory response to insulin resistance or elevated glycaemia. However, we previously found that disrupting the daily glucocorticoid rhythm in mice rapidly increases circulating insulin several-fold while fasting glucose remains normal. This raised the question of what generates and sustains the hyperinsulinaemia. Because glucocorticoids act directly on beta cells through the glucocorticoid receptor (GR), we tested whether beta cell GR is required for this rise in insulin and whether elevated insulin is necessary to maintain glucose homeostasis.

**Methods:** Glucocorticoid rhythms were disrupted in male C57BL/6J mice by subcutaneously implanting corticosterone pellets that raise the trough and lower the peak while maintaining near-physiological mean glucocorticoid exposure, a manipulation we refer to as GC-flattening. Placebo-treated mice served as controls, and high-fat-diet-fed mice provided a metabolic comparison. Beta cell function was assessed by dynamic glucose-stimulated insulin secretion and beta cell–specific Ca²⁺ imaging. The requirement for beta cell GR was tested using adult-inducible beta cell–specific GR knockout mice, thereby limiting developmental effects of constitutive GR deletion. Combined beta cell and hepatocyte GR knockout mice were used to test the consequences of further reducing systemic insulin availability. Insulin sensitivity and glucose tolerance were assessed in vivo. Insulin clearance was assessed from plasma C-peptide:insulin ratios, direct measurement of the disappearance of intravenously administered human insulin, and hepatic insulin-degrading enzyme abundance and activity.

**Results:** GC-flattening produced rapid, sustained hyperinsulinaemia while blood glucose remained normal, distinct from the more gradual hyperinsulinaemia and hyperglycaemia observed in high-fat-diet-fed mice. Islets from GC-flattened mice retained enhanced insulin secretion and Ca²⁺ responses to glucose after isolation, indicating a persistent increase in beta cell glucose responsiveness. During GC-flattening, beta cell GR deletion reduced cumulative circulating insulin exposure by approximately 40% (p < 0.001) and worsened glycaemic control despite similar or greater insulin sensitivity, demonstrating that the GR-dependent rise in insulin helps maintain glucose homeostasis. Direct measurement on Day 3 confirmed reduced insulin clearance in GC-flattened mice. This was accompanied by a reduced plasma C-peptide-to-insulin ratio and decreased hepatic insulin-degrading enzyme abundance and activity. Further lowering circulating insulin by combined beta cell and hepatocyte GR deletion worsened glycaemic control further.

**Conclusions/interpretation:** Disruption of glucocorticoid rhythmicity directly initiates hyperinsulinaemia by enhancing glucose-stimulated insulin secretion through beta cell GR signalling and by reducing insulin clearance. The resulting increase in insulin is required to maintain glucose homeostasis, demonstrating that hyperinsulinaemia can be an early adaptive response to altered endocrine timing rather than simply a consequence of insulin resistance or hyperglycaemia.

**RESEARCH IN CONTEXT:** *What is already known about this subject?:*

- Flattened cortisol rhythms occur in chronic stress, sleep disruption, circadian misalignment, ageing and states of cortisol excess and are associated with adverse metabolic outcomes.
- Glucocorticoid rhythm flattening in mice produces rapid, sustained hyperinsulinaemia while blood glucose remains normal.
- Circulating insulin levels reflect both beta cell secretion and insulin clearance.

*What is the key question?:*

- How does glucocorticoid rhythm disruption increase circulating insulin, and is beta cell glucocorticoid receptor signalling required for the insulin response and for maintaining glucose homeostasis?

*What are the new findings?:*

- GC-flattening increases glucose-stimulated insulin secretion, and beta cell GR is required for much of this enhanced secretory response.
- Adult beta cell GR deletion markedly blunts the rise in circulating insulin and worsens glycaemic control despite similar or greater insulin sensitivity.
- GC-flattening also reduces insulin clearance, providing a second mechanism that increases circulating insulin.

*How might this impact on clinical practice in the foreseeable future?:*

- These findings suggest that cortisol timing, not only overall cortisol exposure, may contribute to metabolic risk and help explain how hyperinsulinaemia and insulin resistance can develop before overt hyperglycaemia.

## INTRODUCTION

Hyperinsulinaemia is one of the earliest and most common endocrine changes in metabolic disease and is generally interpreted as compensation for insulin resistance [1–3]. As insulin sensitivity declines, higher circulating insulin can help maintain glucose homeostasis and thereby delay the development of overt hyperglycaemia. However, circulating insulin is not determined solely by the metabolic demand for insulin. Beta cell secretion is regulated by hormonal, neural and metabolic signals [4–6], and circulating insulin levels are also strongly influenced by insulin clearance. These mechanisms raise the possibility that hyperinsulinaemia can be actively generated or amplified rather than arising solely as a secondary response to insulin resistance or increased glycaemia [6–8].

Our previous studies identified altered glucocorticoid timing as a physiological perturbation that produces a rapid and unusually large increase in circulating insulin. Circulating glucocorticoid levels normally follow a daily rhythm with a pronounced peak and trough [9, 10]. Flattening of this rhythm, in which the normal daily trough is raised and the peak is blunted while mean glucocorticoid levels remain near physiological levels (Fig. 1a), occurs with chronic stress [11, 12], and altered glucocorticoid rhythms are also observed with irregular or insufficient sleep and circadian disruption [13, 14]. In rodents, raising trough corticosterone while maintaining near-physiological mean glucocorticoid exposure produces a rapid, several-fold increase in circulating insulin that persists as substantial insulin resistance develops, while blood glucose remains normal (Fig. 1b, left) [12, 15, 16]. This trajectory differs from HFD feeding (Fig. 1b, right), in which insulin resistance increases insulin demand and hyperglycaemia develops when beta cell compensation becomes insufficient [17–19]. These observations suggested that GC-flattening establishes a distinct hyperinsulinaemic state and raised the question of what mechanisms generate and sustain the increase in circulating insulin.

**Fig. 1.**
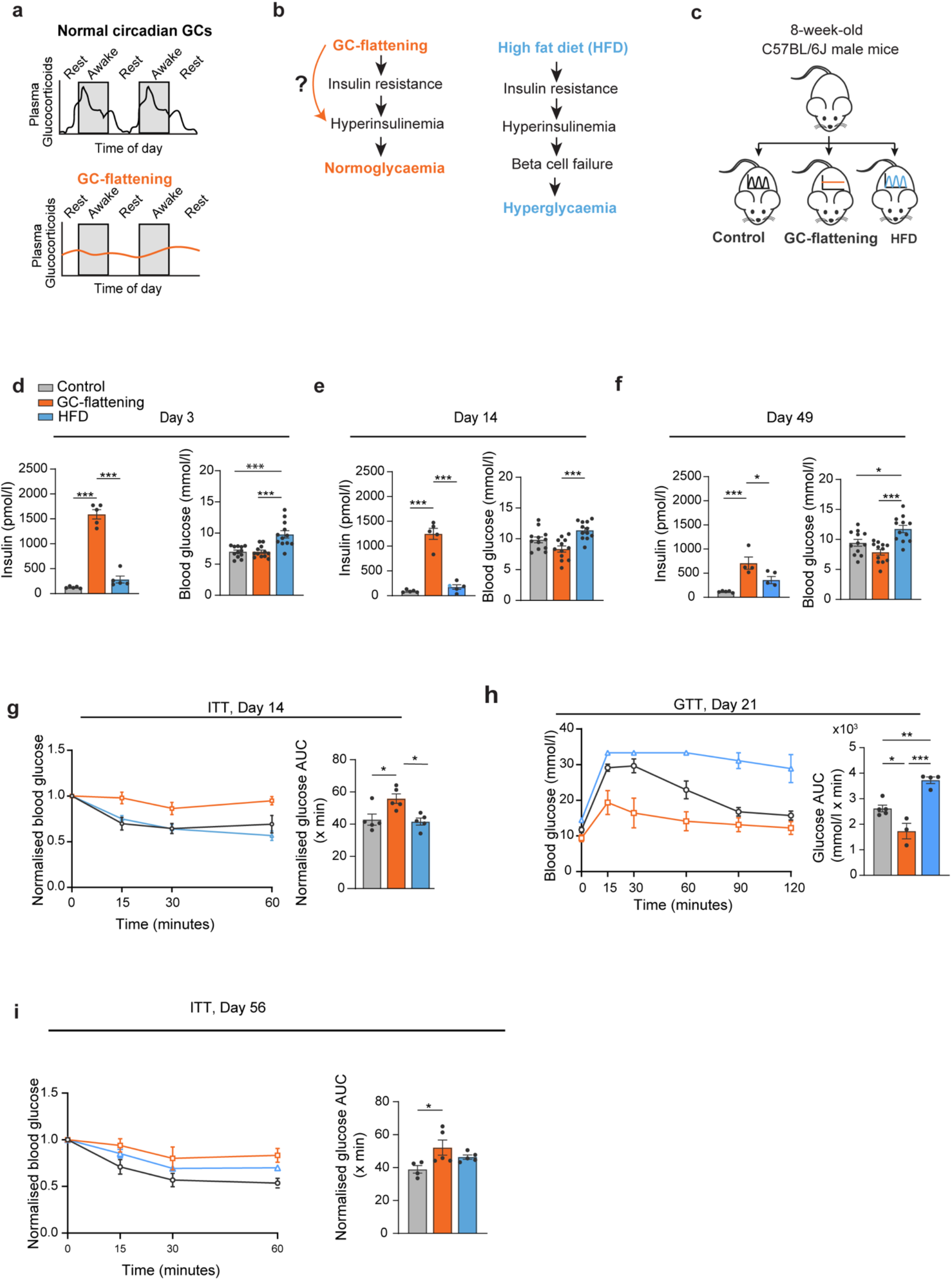
GC-flattening induces sustained hyperinsulinaemia with maintained normoglycaemia, distinct from high-fat diet feeding. (a) Schematic representation of normal circadian and flattened plasma glucocorticoid rhythms. GC-flattening raises trough glucocorticoid levels and reduces peak amplitude. (b) Conceptual models contrasting the metabolic progression during high-fat diet (HFD) feeding with the state induced by GC-flattening. (c) Experimental design. Eight-week-old male C57BL/6J mice received placebo pellets while maintained on chow (Control), corticosterone pellets while maintained on chow (GC-flattening), or placebo pellets while fed HFD (HFD). The question mark in (b) indicates the unresolved mechanism by which GC-flattening produces a rapid and sustained increase in circulating insulin beyond the classical compensatory response to insulin resistance. (d–f) Fasting plasma insulin and blood glucose at Day 3 (d), Day 14 (e) and Day 49 (f). GC-flattened mice developed sustained hyperinsulinaemia from Day 3 while maintaining normoglycaemia. HFD-fed mice showed a smaller, later increase in fasting insulin and progressive elevations in fasting glucose. For insulin, n=5 mice per group except GC-flattening at Day 49, n=4. For glucose, n=12 mice per group. (g) Insulin tolerance test (ITT) at Day 14, showing normalised blood glucose after intraperitoneal insulin administration (0.5 U/kg) and corresponding area under the curve (AUC) calculated from the normalised glucose values. GC-flattened mice were insulin resistant, whereas HFD-fed mice remained as insulin sensitive as controls. n=5 mice per group. (h) Glucose tolerance test (GTT) at Day 21 after intraperitoneal glucose administration (2 g/kg) and corresponding AUC. Despite insulin resistance, GC-flattened mice showed improved glucose tolerance compared with controls, whereas HFD-fed mice were glucose intolerant. n=5 Control, n=3 GC-flattened and n=4 HFD mice. (i) ITT at Day 56 and corresponding AUC calculated from the normalised glucose values. By this later timepoint, both GC-flattened and HFD-fed mice were insulin resistant relative to controls, with no significant difference between the two treatment groups. n=7 mice per group. Data are mean ± SEM. Group comparisons shown in (d–i) were performed by one-way ANOVA followed by Tukey’s multiple-comparisons test. Blood glucose values reaching the glucometer limit were recorded as 33.3 mmol/l, making AUC values for animals with censored measurements underestimates. Asterisks indicate Tukey-adjusted multiple-comparison p values: *p<0.05, **p<0.01 and ***p<0.001. Only statistically significant pairwise comparisons are shown in the panels.

The pancreatic beta cell is a plausible target through which altered glucocorticoid timing could increase circulating insulin because beta cells express the glucocorticoid receptor (GR). Most previous studies of glucocorticoid action on beta cells have used sustained or pharmacological glucocorticoid exposure, which is generally associated with impaired beta cell function and reduced glucose-stimulated insulin secretion [20, 21]. However, in vivo glucocorticoid treatment in rats and humans can instead increase insulin secretion, and studies in both species have reported enhanced beta cell glucose responsiveness [22–25]. These different outcomes suggest that glucocorticoid effects on beta cell secretion depend on the dose, duration and temporal pattern of exposure. Whether disruption of the normal glucocorticoid rhythm increases beta cell glucose responsiveness through GR, and whether beta cell GR signalling is required for the rise in circulating insulin during GC-flattening, have not been established.

This question is clinically relevant because altered cortisol rhythms are observed with chronic stress, sleep disruption and ageing [11–14, 26, 27], as well as in Cushing’s syndrome [28], and mild autonomous cortisol secretion (MACS) [29]. A flatter diurnal cortisol slope predicts future impaired fasting glucose or type 2 diabetes [30] and is associated with increased all-cause and cardiovascular mortality [31]. Among middle-aged men, stress-related cortisol secretion in the setting of low diurnal cortisol variation was strongly associated with abdominal obesity, fasting insulin and blood pressure [32]. GC-flattening isolates one feature of cortisol-rhythm disruption—the loss of a pronounced daily trough—while maintaining mean glucocorticoid exposure within the physiological range [11, 12, 15, 16, 33, 34]. It therefore allows us to test whether loss of the daily glucocorticoid trough itself alters insulin regulation.

Here we asked how disrupted glucocorticoid timing raises circulating insulin, and whether the resulting hyperinsulinaemia is required to maintain glucose homeostasis. Because circulating insulin reflects both beta cell secretion and insulin clearance, much of which occurs in the liver, we examined each in turn. We assessed beta cell glucose responsiveness by islet perifusion and beta cell Ca²⁺ imaging, and tested whether beta cell GR is required using adult-inducible beta cell-specific deletion. We then measured insulin clearance directly, and used combined beta cell and hepatocyte GR deletion to lower circulating insulin further and assess the consequences for glycaemic control.

## METHODS

### Animals and glucocorticoid-rhythm flattening

Male mice aged 8–14 weeks at pellet implantation were used unless otherwise stated and housed under a 12-h light/dark cycle with standard chow. Wild-type mice were C57BL/6J mice obtained from The Jackson Laboratory (stock #000664), and genetically modified lines were maintained on a C57BL/6J background. A high-fat diet providing 60% of energy from fat was used for diet-induced obesity studies. Glucocorticoid rhythms were flattened by subcutaneous corticosterone pellets delivering approximately 10 mg kg⁻¹ day⁻¹, which raise trough levels and reduce daily peaks while maintaining mean daily corticosterone exposure within the physiological range [15, 16, 33, 34]. Placebo pellets served as controls. Male mice were used to maintain comparability with the cohorts in which the GC-flattening phenotype was originally characterised [15, 16, 33]. All animal procedures were approved by the Institutional Animal Care and Use Committee of Weill Cornell Medical College (protocol number 2020-0001), and experiments were performed in accordance with the NIH Guide for the Care and Use of Laboratory Animals and applicable institutional and federal regulations.

### Mouse models

Beta cell–specific GR knockout (beta cell GRKO) mice were generated by crossing MIP-CreERT (The Jackson Laboratory, stock #024709) [35] with *Nr3c1^fl/fl^* (GRfl/fl) mice (stock #021021) [36]. Cre-negative GRfl/fl and MIP-CreERT littermates served as controls.

Recombination was induced in adult mice (8–10 weeks) with tamoxifen (75 mg/kg intraperitoneally once daily for 5 days). Double-GRKO mice, with combined beta cell and hepatocyte GR deletion, were generated in a separate cohort by retro-orbital injection of AAV8-TBG-iCre (Vector Biolabs, VB1724) into beta cell GRKO mice. The MIP-CreERT and beta cell GRKO groups shown in Fig. 6 did not receive AAV. Hepatic GR deletion in the double-GRKO group was induced with AAV8-TBG-iCre, with AAV8-TBG-LacZ used in separate AAV control experiments. Beta cell Ca²⁺ responses were measured in Ins1-Cre;GCaMP6f reporter mice, generated by crossing Ins1-Cre mice (stock #026801) [37] with a Cre-dependent GCaMP6f reporter line (stock #028865) [38] so that the calcium indicator was expressed specifically in beta cells.

### In vivo phenotyping

After a 5-h fast, blood glucose was measured with a glucometer and plasma insulin and C-peptide by immunoassay. Insulin tolerance tests (0.5 U/kg human insulin) were performed with blood sampling at 0, 15, 30 and 60 min, and glucose tolerance tests (2 g/kg glucose) with sampling at 0, 15, 30, 60, 90 and 120 min.

### Ex vivo assays

Islets were isolated by collagenase digestion. Dynamic glucose-stimulated insulin secretion was measured by perifusion (3 → 8 → 3 → 20 mmol/l glucose) and expressed as pmol/l per 100 islet equivalents. Islet insulin content was determined by acid–ethanol extraction and insulin immunoassay (ESM Methods). Beta cell–specific GCaMP6f Ca²⁺ dynamics were imaged in intact islets from Ins1-Cre;GCaMP6f mice across the same glucose steps, with fluorescence quantified from whole-islet regions of interest. Hepatic IDE and CEACAM1 abundance were assessed by immunoblotting (vinculin loading control) and IDE activity by a fluorometric assay. GR-deletion efficiency was confirmed by flow cytometry and immunofluorescence (beta cells, ESM Fig. 3). Hepatocyte GR deletion was confirmed in the companion study [16].

### Statistical analysis

Data are presented as mean ± SEM. For in vivo experiments, the experimental unit was the individual mouse. For ex vivo assays, n denotes independent biological replicates as defined in the corresponding figure legends and ESM; individual islets contributing to averaged Ca²⁺ traces were not treated as independent biological replicates. Statistical tests and exact n values are specified in the corresponding figure legends. A value of p<0.05 was considered statistically significant. Analyses were performed in GraphPad Prism v11 (GraphPad Software, San Diego, CA, USA).

Full protocols, reagents, antibodies and image-acquisition settings are provided in the electronic supplementary material (ESM) Methods.

## RESULTS

### GC-flattening produces a hyperinsulinaemic, normoglycaemic state

GC-flattening produces a rapid, sustained rise in circulating insulin while fasting glucose remains normal [15, 16], raising the question of how disrupted glucocorticoid timing increases insulin without an accompanying rise in glycaemia. Figure 1a illustrates the loss of the normal daily glucocorticoid trough produced by GC-flattening. We contrasted the resulting metabolic state with the canonical progression during high-fat-diet (HFD) feeding (Fig. 1b). During GC-flattening, insulin resistance develops, but circulating insulin rises rapidly to high levels while fasting glucose remains normal (Fig. 1b, left). In HFD-fed mice, by contrast, progressive insulin resistance drives compensatory hyperinsulinaemia, followed by loss of glycaemic control as compensation becomes insufficient [18, 19] (Fig. 1b, right). We therefore considered that the hyperinsulinaemia induced by GC-flattening may involve mechanisms beyond the classical HFD-like response to insulin resistance.

To test whether these two conditions follow different metabolic trajectories, we compared fasting insulin and glucose over time in control, GC-flattened and HFD-fed mice (Fig. 1c–f). By Day 3, GC-flattened mice exhibited a several-fold increase in fasting insulin, whereas insulin in HFD-fed mice remained comparable to controls (Fig. 1d). Despite this marked increase in insulin, fasting glucose remained normal in GC-flattened mice, while HFD-fed mice already exhibited significantly higher glucose. At Day 14, hyperinsulinaemia remained pronounced in GC-flattened mice, whereas insulin in HFD-fed mice remained much lower (Fig. 1e). Fasting glucose was again significantly lower in GC-flattened than in HFD-fed mice. By Day 49, fasting insulin remained significantly elevated in GC-flattened mice and was also higher than in HFD-fed mice, while the increase in HFD-fed mice did not reach significance relative to controls (Fig. 1f). At this later timepoint, fasting glucose remained normal in GC-flattened mice but was significantly elevated in HFD-fed mice. Thus, GC-flattening produces a rapid and sustained increase in circulating insulin while maintaining normoglycaemia, in contrast to the slower insulin response and progressive hyperglycaemia during HFD feeding.

We also compared insulin sensitivity and glucose tolerance between the two conditions. At Day 14, GC-flattened mice were already insulin resistant, whereas HFD-fed mice remained as insulin sensitive as controls (Fig. 1g). Thus, insulin resistance develops rapidly during GC-flattening, before a comparable defect is detectable during HFD feeding.

Despite this early insulin resistance, GC-flattened mice showed significantly improved glucose tolerance at Day 21 compared with controls, whereas HFD-fed mice were glucose intolerant (Fig. 1h). Thus, GC-flattened mice maintain improved glucose tolerance despite impaired insulin sensitivity. This striking dissociation between insulin sensitivity and glucose tolerance distinguishes the metabolic state induced by GC-flattening from that produced by HFD.

To determine whether the difference in insulin sensitivity between GC-flattened and HFD-fed mice reflected distinct severity or different kinetics, we repeated the ITT at Day 56 (Fig. 1i). By this later timepoint, both GC-flattened and HFD-fed mice were insulin resistant relative to controls, and insulin sensitivity no longer differed significantly between the two treatment groups. Thus, GC-flattening induces insulin resistance substantially earlier than HFD feeding, although both groups were insulin resistant by Day 56 and no longer differed significantly from one another. This earlier onset is consistent with the companion study, in which molecular defects in skeletal muscle insulin signalling were detectable within days of GC-flattening[16]. Because circulating insulin increased markedly while glycaemia remained normal, the hyperinsulinaemia could not be explained simply by increased glycaemic demand. Moreover, GC-flattened mice showed improved glucose tolerance despite impaired insulin sensitivity. We therefore asked whether GC-flattening directly alters beta cell insulin secretion.

### GC-flattening establishes a persistent increase in glucose-stimulated insulin secretion

The rapid rise in circulating insulin during GC-flattening occurs without an accompanying increase in fasting glucose (Fig. 1), raising the possibility that glucocorticoids directly increase beta cell secretory responsiveness. To test for a direct effect, we isolated islets from wild-type mice, treated them with corticosterone (280 nmol/l) or vehicle for 24 h and measured dynamic insulin secretion during sequential perifusion with 3, 8, 3 and 20 mmol/l glucose (Fig. 2a).

**Fig. 2.**
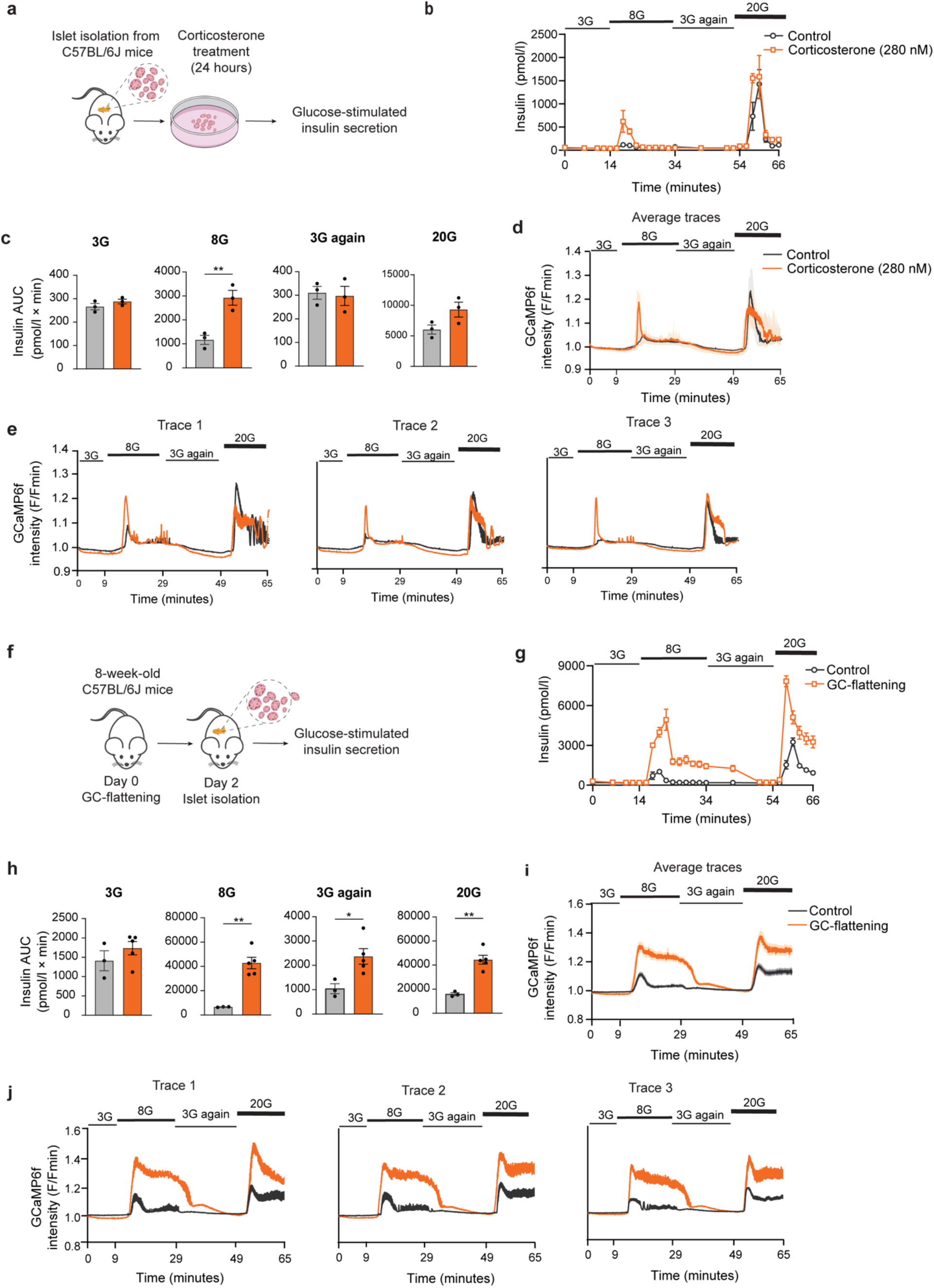
GC-flattening establishes a persistent increase in beta cell glucose responsiveness. (a) Experimental design for treatment of isolated C57BL/6J islets with corticosterone (280 nmol/l) or vehicle for 24 h before dynamic glucose-stimulated insulin secretion (GSIS). (b) Dynamic insulin secretion during sequential perifusion with 3, 8, 3 and 20 mmol/l glucose. (c) AUC for each glucose phase in (b). Corticosterone increased insulin secretion at 8 mmol/l glucose, with no significant difference during the initial 3 mmol/l phase, after return to 3 mmol/l glucose or during the 20 mmol/l phase. (d, e) GCaMP6f Ca²⁺ responses in islets from Ins1-Cre;GCaMP6f mice treated with corticosterone or vehicle during the same glucose sequence. Average traces (d) and representative individual islet traces (e) are shown. Corticosterone-treated islets exhibited enhanced glucose-triggered Ca²⁺ responses. (f) Experimental design for GC-flattening in vivo followed by islet isolation on Day 2 and dynamic GSIS. (g) Dynamic insulin secretion from islets isolated from control and GC-flattened mice during sequential perifusion with 3, 8, 3 and 20 mmol/l glucose. (h) AUC for each glucose phase in (g). GC-flattening increased secretion during the 8 mmol/l phase, after return to 3 mmol/l glucose and during the 20 mmol/l phase. (i, j) GCaMP6f Ca²⁺ responses in islets isolated from control and GC-flattened mice. Average traces (i) and representative individual islet traces (j) are shown. Ca²⁺ responses were enhanced during 8 mmol/l glucose, remained elevated after return to 3 mmol/l glucose and were increased during subsequent 20 mmol/l glucose stimulation. For GSIS experiments, n=3 independent biological replicates per group. For the in vitro corticosterone experiment in (b, c), islets from each biological replicate were pooled and divided between vehicle and corticosterone treatment, and AUC values for each glucose phase were compared using paired two-tailed t tests. For the in vivo GC-flattening experiment in (g, h), control and GC-flattened mice were independent groups, and AUC values for each glucose phase were compared using unpaired two-tailed t tests. For Ca²⁺ imaging in (d, e), n=3 independent biological replicates, each from a separate islet isolation, with islets from each isolation divided between vehicle and corticosterone treatment. For Ca²⁺ imaging in (i, j), n=2 independent biological replicates per group, with one placebo-treated and one GC-flattened mouse analysed in each replicate. Each average Ca²⁺ trace comprised 5–15 islets. Data are mean ± SEM. Asterisks indicate p values: *p<0.05 and **p<0.01. The Ca²⁺ imaging conditions are detailed in the ESM Methods. Only statistically significant pairwise comparisons are shown on the panels.

Corticosterone-treated islets showed increased insulin secretion during stimulation with 8 mmol/l glucose (Fig. 2b). Quantification of the integrated secretory response confirmed a significant increase at 8 mmol/l glucose, whereas secretion during the initial 3 mmol/l phase, after return to 3 mmol/l glucose and during 20 mmol/l glucose did not differ significantly between groups (Fig. 2c). Thus, direct corticosterone exposure increases insulin secretion in response to 8 mmol/l glucose.

To determine whether this increased secretory response was accompanied by altered glucose-triggered beta cell Ca²⁺ signalling, we imaged Ca²⁺ in intact islets from Ins1-Cre;GCaMP6f reporter mice, in which GCaMP6f is expressed specifically in beta cells. Corticosterone-treated islets showed an enhanced Ca²⁺ response following the transition from 3 to 8 mmol/l glucose, together with altered and more sustained Ca²⁺ responses during stimulation with 20 mmol/l glucose (Fig. 2d, e; ESM Fig. 2). These findings demonstrate that the effect of corticosterone on insulin secretion is accompanied by increased glucose-triggered beta cell activation.

We therefore tested whether a similar secretory state develops during GC-flattening and persists after islet isolation. Islets were isolated two days after placebo or corticosterone pellet implantation and subsequently analysed during sequential stimulation with 3, 8, 3 and 20 mmol/l glucose (Fig. 2f). Islets from GC-flattened mice exhibited a marked increase in insulin secretion during stimulation with 8 mmol/l glucose and a larger response to 20 mmol/l glucose than islets from circadian controls (Fig. 2g). Unlike the 24-h corticosterone treatment, the effect established *in vivo* also persisted after glucose was returned from 8 to 3 mmol/l. Quantification confirmed significantly increased secretion during the 8 mmol/l, second 3 mmol/l and 20 mmol/l phases, whereas secretion during the initial 3 mmol/l glucose phase was unchanged (Fig. 2h). Thus, GC-flattening establishes a broader and more persistent change in beta cell secretory responsiveness than corticosterone treatment of isolated islets alone.

Consistent with the secretion data, GCaMP6f imaging revealed a pronounced increase in glucose-triggered Ca²⁺ signalling in islets from GC-flattened mice (Fig. 2i, j). Ca²⁺ responses were enhanced during stimulation with 8 mmol/l glucose, remained elevated after glucose was returned to 3 mmol/l and showed a larger sustained response during subsequent stimulation with 20 mmol/l glucose. The persistence of both the secretory and Ca²⁺ phenotypes after islet isolation indicates that GC-flattening establishes a beta cell state that does not require the continued presence of the *in vivo* hormonal or glycaemic environment.

Because increased secretion could potentially deplete stored insulin, we also measured total islet insulin content after static glucose stimulation in islets isolated after 14 days of GC-flattening. Insulin content was not reduced in GC-flattened islets after high-glucose or KCl stimulation and was significantly increased under low-glucose conditions (ESM Fig. 1). Thus, the enhanced secretion induced by GC-flattening occurs without depletion of islet insulin stores. Together, these experiments show that glucocorticoid exposure directly enhances glucose-stimulated insulin secretion and that GC-flattening establishes a stronger secretory phenotype that persists after islet isolation.

### Beta cell glucocorticoid receptor signalling drives the insulin rise during GC-flattening

Having established that glucocorticoid exposure enhances glucose-stimulated insulin secretion, we tested whether beta cell glucocorticoid receptor (GR) signalling is required for the rise in circulating insulin during GC-flattening. We generated mice with beta cell–specific deletion of GR induced in adulthood using a tamoxifen-inducible MIP-CreERT;GRfl/fl system (Fig. 3a, b). This strategy allowed us to test the function of GR in mature beta cells while limiting developmental adaptation as an explanation for the phenotype. Flow cytometry confirmed efficient deletion, with approximately 89% of insulin-positive cells lacking detectable GR, and immunofluorescence of isolated islets independently confirmed loss of beta cell GR (ESM Fig. 3).

**Fig. 3.**
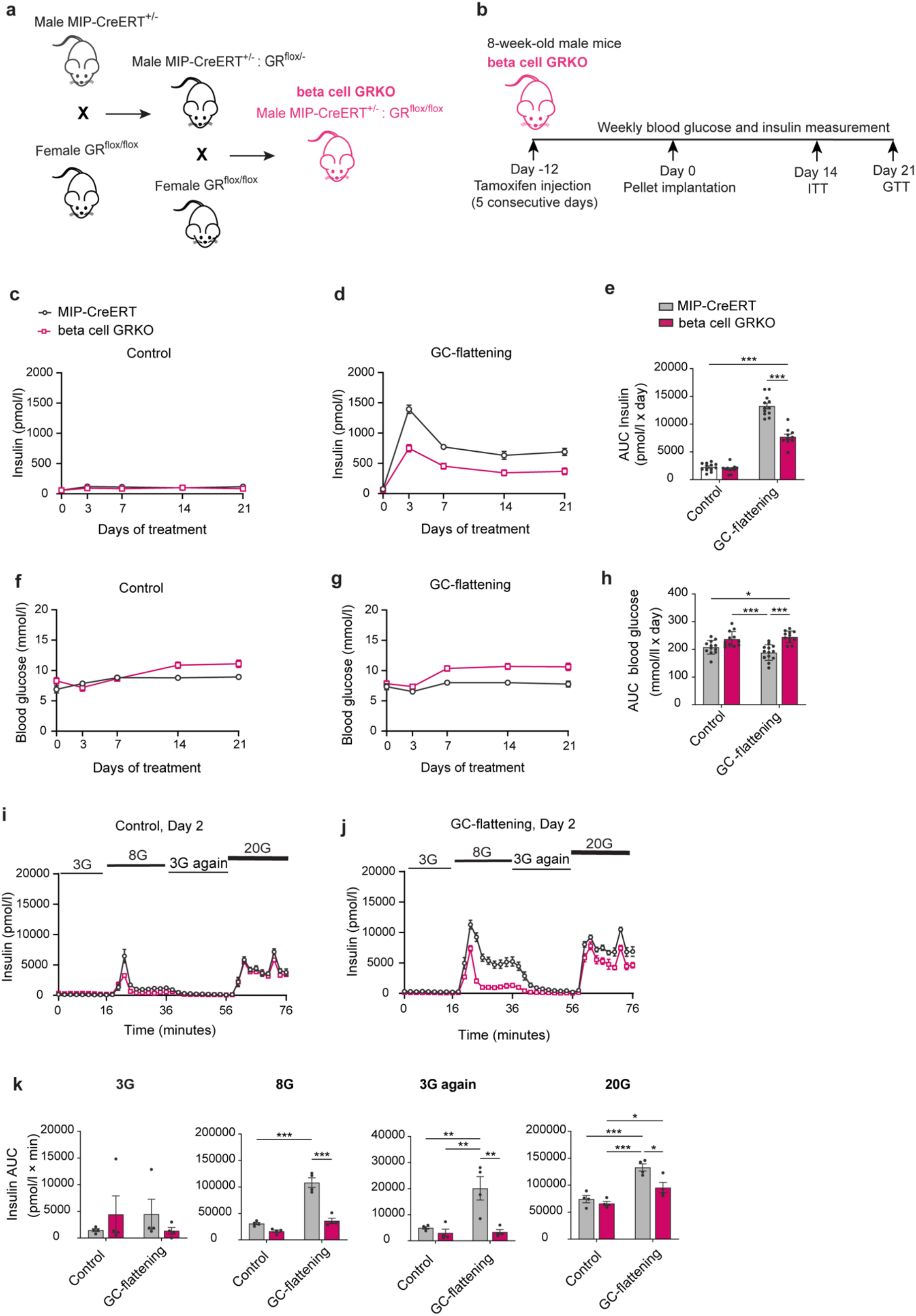
Loss of beta cell GR blunts GC-flattening-induced hyperinsulinaemia and causes hyperglycaemia. (a) Breeding scheme for generation of adult-inducible beta cell-specific glucocorticoid receptor knockout (beta cell GRKO) mice using MIP-CreERT;GR^fl/fl mice. (b) Experimental timeline showing tamoxifen induction, placebo or corticosterone pellet implantation and metabolic testing. (c, d) Fasting plasma insulin over time in MIP-CreERT control and beta cell GRKO mice under placebo-pellet control (c) and GC-flattened (d) conditions. (e) AUC for circulating insulin in (c, d). GC-flattening increased insulin in both genotypes, but the increase was markedly reduced in beta cell GRKO mice. n=12, 11, 12 and 10 for MIP-CreERT control, beta cell GRKO control, MIP-CreERT GC-flattened and beta cell GRKO GC-flattened groups, respectively. (f, g) Fasting blood glucose over time under placebo-pellet control (f) and GC-flattened (g) conditions. Beta cell GRKO mice had higher glucose under both conditions, with a larger difference during GC-flattening. (h) Glucose AUC for (f, g). n=12, 12, 13 and 12 for the corresponding groups. (i, j) Dynamic GSIS from islets isolated on Day 2 after placebo (i) or corticosterone (j) pellet implantation. (k) Insulin AUC for the initial 3 mmol/l, 8 mmol/l, second 3 mmol/l and 20 mmol/l glucose phases. GC-flattening increased secretion in MIP-CreERT islets during the 8 mmol/l, second 3 mmol/l and 20 mmol/l phases. No significant GC-flattening-induced increase was detected in beta cell GRKO islets at 8 mmol/l or during the second 3 mmol/l phase. At 20 mmol/l glucose, secretion also increased in beta cell GRKO islets but remained significantly lower than in MIP-CreERT islets. Under GC-flattened conditions, secretion was significantly lower in beta cell GRKO than in MIP-CreERT islets during the 8 mmol/l, second 3 mmol/l and 20 mmol/l phases. n=4 mice per group. Data are mean ± SEM. AUC measurements were analysed by two-way ANOVA with Tukey’s multiple-comparisons test. For secretion at 8 mmol/l glucose, the treatment × genotype interaction was significant (F(1,12)=27.9, p<0.001). Each glucose phase was analysed independently. Asterisks indicate Tukey-adjusted multiple-comparison p values: *p<0.05, **p<0.01 and ***p<0.001. Only statistically significant pairwise comparisons are shown on the panels.

Following recombination in adult mice, beta cell GRKO mice and MIP-CreERT littermate controls received placebo or corticosterone pellets. Under placebo conditions, circulating insulin remained low and comparable between genotypes throughout the observation period (Fig. 3c). In contrast, GC-flattening produced a rapid and sustained increase in circulating insulin in MIP-CreERT controls, whereas this response was markedly attenuated in beta cell GRKO mice (Fig. 3d). Quantification of cumulative insulin exposure confirmed that GC-flattening strongly increased insulin AUC in MIP-CreERT controls and that cumulative insulin exposure during GC-flattening was approximately 40% lower in beta cell GRKO mice than in MIP-CreERT controls (44.8 ± 2.8 vs 77.0 ± 3.0; p < 0.001; Fig. 3e). Thus, beta cell GR signalling is a major contributor to the rise in circulating insulin induced by GC-flattening. However, GC-flattening still increased circulating insulin in beta cell GRKO mice, indicating that beta cell GR accounts for only part of the hyperinsulinaemic response.

Loss of beta cell GR also had a clear glycaemic consequence. Beta cell GRKO mice had higher fasting glucose than MIP-CreERT controls under both placebo and GC-flattened conditions, with the difference becoming substantially greater during GC-flattening (Fig. 3f, g). Under GC-flattened conditions, glucose in MIP-CreERT controls remained relatively stable, whereas glucose in beta cell GRKO mice increased from Day 7 onwards (Fig. 3g). Cumulative glucose exposure was significantly higher in beta cell GRKO mice under both conditions, with the greatest increase during GC-flattening (Fig. 3h). These findings indicate that beta cell GR contributes to glycaemic control under control conditions but becomes particularly important when glucocorticoid rhythmicity is disrupted.

To determine whether the reduced circulating insulin in beta cell GRKO mice reflected loss of the enhanced beta cell secretory phenotype established during GC-flattening, we isolated islets from MIP-CreERT control and beta cell GRKO mice two days after placebo or corticosterone pellet implantation and analysed them by dynamic glucose perifusion (Fig. 3i–k). Under placebo conditions, insulin secretion was comparable between genotypes throughout the glucose-stimulation sequence (Fig. 3i, k). Following GC-flattening, MIP-CreERT islets showed increased secretion at 8 mmol/l glucose, after return to 3 mmol/l glucose and at 20 mmol/l glucose. In beta cell GRKO islets, no significant GC-flattening-induced increase was detected at 8 mmol/l or after return to 3 mmol/l glucose, whereas secretion still increased at 20 mmol/l glucose (Fig. 3j, k). Under GC-flattened conditions, however, secretion was lower in beta cell GRKO than in MIP-CreERT islets at 8 mmol/l glucose, after return to 3 mmol/l glucose and at 20 mmol/l glucose. Thus, beta cell GR is required for much of the enhanced insulin secretion established by GC-flattening. The preserved increase in secretion at 20 mmol/l glucose argues against a general defect in insulin secretion in beta cell GRKO islets.

### Beta cell GR signalling supports glucose tolerance during GC-flattening by sustaining insulin secretion

To distinguish reduced insulin output from worsened peripheral insulin resistance, we measured insulin sensitivity in MIP-CreERT control and beta cell GRKO mice with either normal glucocorticoid rhythmicity or GC-flattening. Under placebo-pellet control conditions, the two genotypes showed similar responses to insulin administration (Fig. 4a, c). GC-flattening significantly reduced insulin sensitivity in MIP-CreERT controls, whereas insulin sensitivity was not significantly altered by GC-flattening in beta cell GRKO mice (Fig. 4b, c). Moreover, under GC-flattened conditions, beta cell GRKO mice showed greater glucose lowering in response to insulin than MIP-CreERT controls (Fig. 4b, c). Thus, the elevated glucose observed after beta cell GR deletion during GC-flattening cannot be attributed to worsened peripheral insulin resistance.

**Fig. 4.**
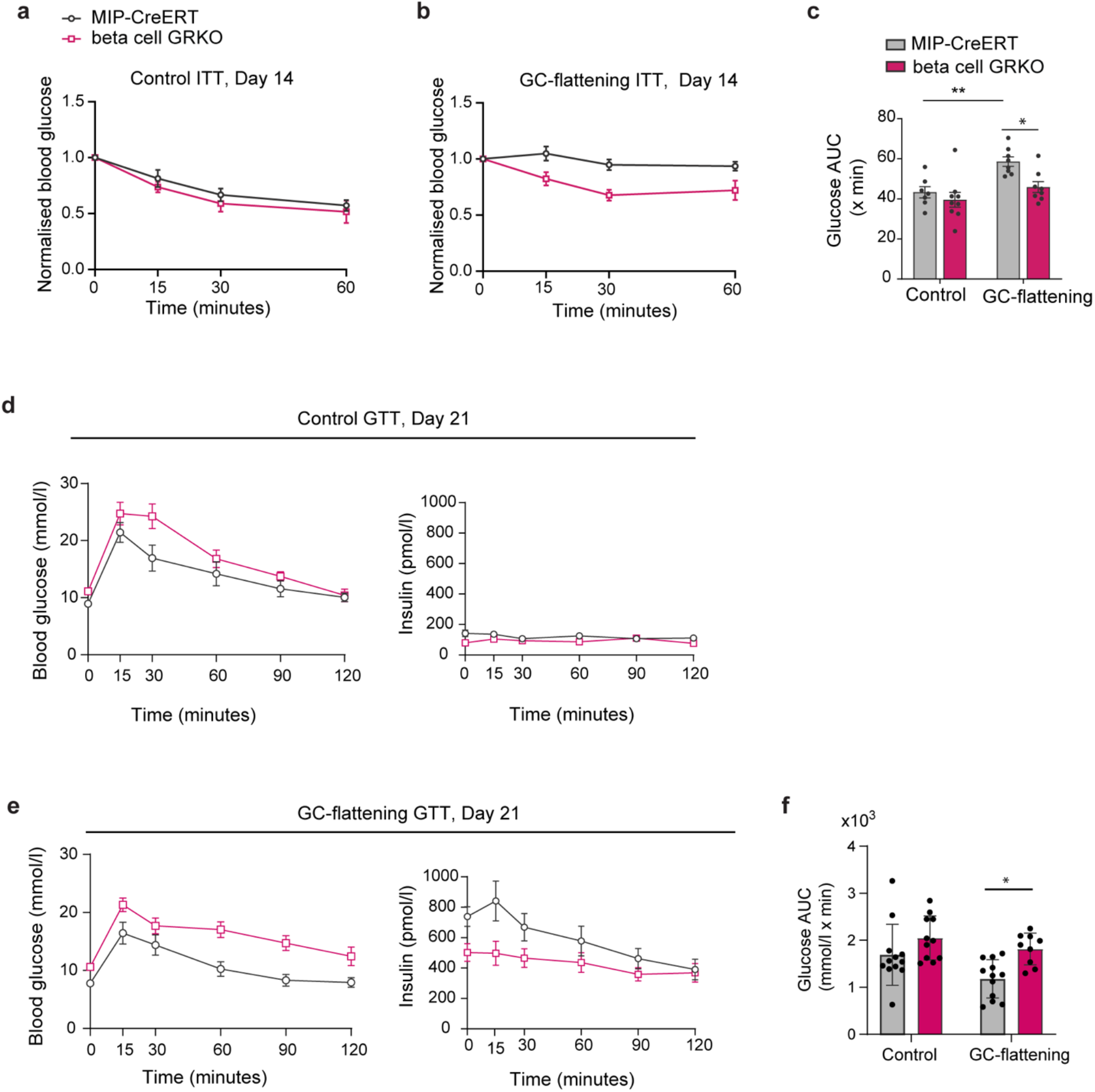
Beta cell GR signalling supports glucose tolerance during GC-flattening by sustaining insulin secretion. (a, b) ITTs at Day 14 under placebo-pellet control (a) and GC-flattened (b) conditions. Blood glucose is expressed relative to baseline. MIP-CreERT and beta cell GRKO mice responded similarly under control conditions, whereas beta cell GRKO mice showed greater insulin-stimulated glucose lowering during GC-flattening. (c) ITT AUC. Under control conditions, ITT AUC did not differ between MIP-CreERT and beta cell GRKO mice (p = 0.861). GC-flattening increased ITT AUC in MIP-CreERT mice (p = 0.007), consistent with reduced insulin sensitivity, but did not significantly alter ITT AUC in beta cell GRKO mice (p = 0.446). Under GC-flattened conditions, ITT AUC was lower in beta cell GRKO mice than in MIP-CreERT controls (p = 0.023), indicating greater insulin sensitivity. ITT AUC was analysed by two-way ANOVA followed by Šídák-adjusted multiple comparisons of four prespecified comparisons. For ITTs, n=7 MIP-CreERT and n=9 beta cell GRKO mice under control conditions and n=8 per genotype under GC-flattened conditions. (d, e) GTTs at Day 21 under placebo-pellet control (d) and GC-flattened (e) conditions. Blood glucose and circulating insulin during the glucose challenge are shown. Under control conditions, glucose excursions were similar overall between genotypes. During GC-flattening, beta cell GRKO mice showed higher glucose excursions and lower circulating insulin than MIP-CreERT controls. (f) Glucose AUC for the GTTs in (d, e). Two-way ANOVA showed a significant main effect of genotype (p = 0.006), with no significant main effect of treatment (p = 0.157) and no genotype × treatment interaction (p = 0.556). Šídák-adjusted multiple comparisons showed no significant difference between MIP-CreERT and beta cell GRKO mice under control conditions (p = 0.208), whereas glucose AUC was higher in beta cell GRKO mice under GC-flattened conditions (p = 0.040). For GTTs, n=12 MIP-CreERT and n=11 beta cell GRKO mice under control conditions and n=12 MIP-CreERT and n=10 beta cell GRKO mice under GC-flattened conditions. For circulating insulin during the GTT, n at 0, 15, 30, 60, 90 and 120 min, respectively, was 8, 9, 11, 12, 11 and 12 for MIP-CreERT mice under control conditions, 12, 12, 12, 11, 11 and 10 for beta cell GRKO mice under control conditions, 10, 10, 10, 9, 10 and 10 for MIP-CreERT mice under GC-flattened conditions, and 11, 9, 10, 11, 10 and 9 for beta cell GRKO mice under GC-flattened conditions. Data are mean ± SEM. Asterisks indicate Šídák-adjusted multiple-comparison p values: *p<0.05 and **p<0.01. Exact p values are reported above. Only statistically significant pairwise comparisons are shown on the panels.

We also examined glucose tolerance and the accompanying insulin response. With normal glucocorticoid rhythmicity, MIP-CreERT and beta cell GRKO mice showed similar overall glucose tolerance (Fig. 4d). Circulating insulin during the glucose challenge was also similar between genotypes. In contrast, during GC-flattening, beta cell GRKO mice exhibited higher glucose excursions and a lower circulating insulin response than MIP-CreERT controls (Fig. 4e). Combined glucose AUC analysis showed a significant genotype effect. Glucose AUC did not differ significantly between genotypes under control conditions but was significantly higher in beta cell GRKO mice during GC-flattening (Fig. 4f).

Beta cell GRKO mice therefore showed worse glucose tolerance during GC-flattening despite greater insulin sensitivity and a smaller insulin response to glucose. Thus, the beta cell GR-dependent insulin response is required to maintain glucose tolerance in this insulin-resistant state.

### Reduced insulin clearance contributes to elevated circulating insulin during GC-flattening

Because beta cell GR deletion markedly reduced but did not abolish the hyperinsulinaemic response to GC-flattening, we asked what additional mechanism contributes to the elevation in circulating insulin. Circulating insulin reflects the balance between beta cell secretion and insulin clearance [39], so we tested whether reduced insulin clearance provides a second mechanism for increasing systemic insulin availability. After 14 days of GC-flattening, both plasma C-peptide and insulin were significantly increased relative to controls (Fig. 5a). The increase in C-peptide is consistent with increased beta cell secretion. However, the C-peptide-to-insulin molar ratio was significantly reduced (p=0.006), indicating that the increase in circulating insulin was disproportionate to the increase in C-peptide (Fig. 5a). Consistent with this interpretation, the relationship between insulin and C-peptide was shifted in GC-flattened mice, with substantially higher circulating insulin for a given C-peptide concentration than in controls (Fig. 5b). Together, these measurements suggested that increased secretion is accompanied by reduced insulin clearance.

**Fig. 5.**
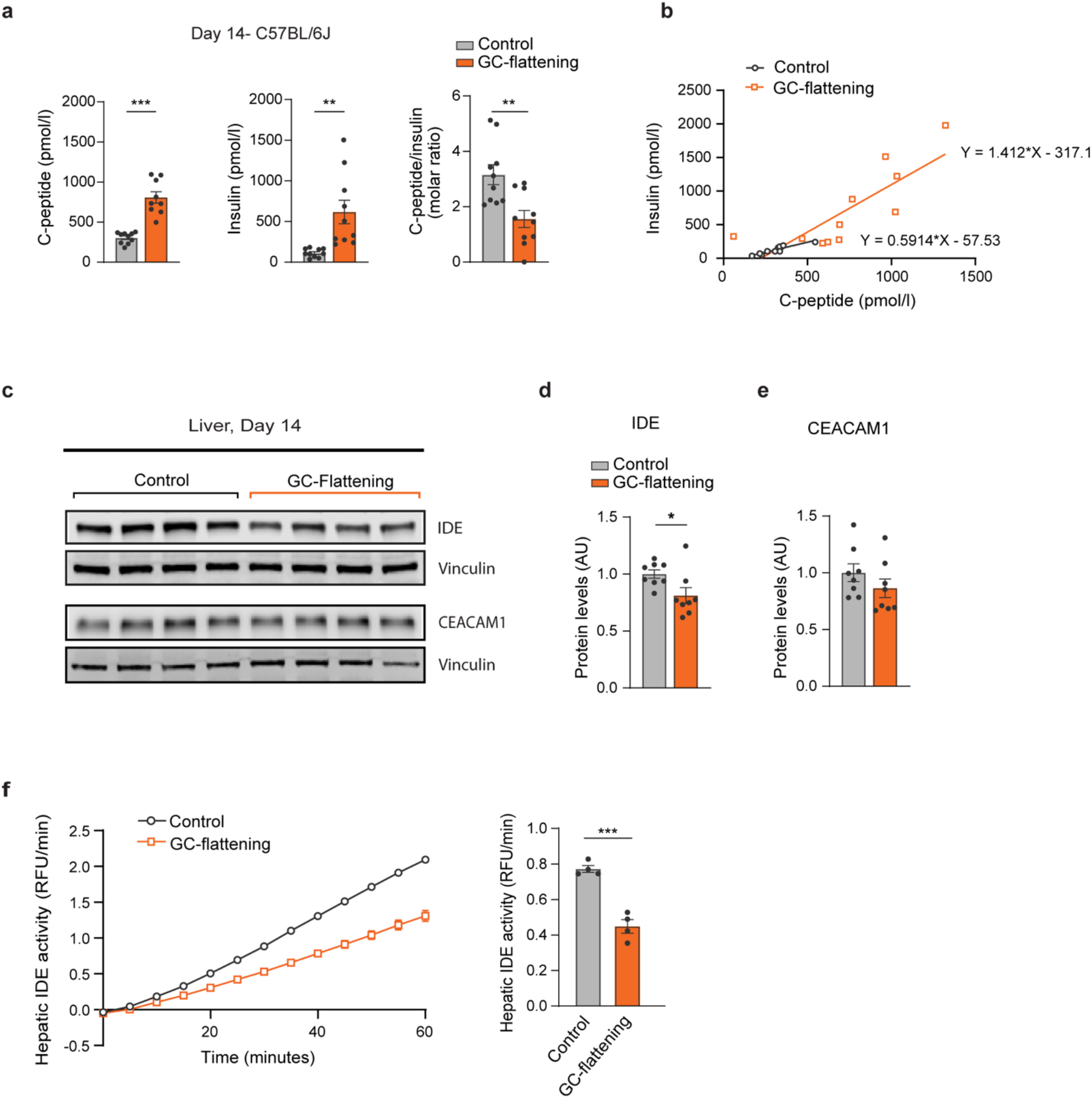
Reduced insulin clearance during GC-flattening is associated with decreased hepatic IDE abundance and activity. (a) Plasma C-peptide, insulin and C-peptide-to-insulin molar ratio after 14 days of control or GC-flattening treatment in C57BL/6J mice. GC-flattening increased C-peptide and insulin and reduced the C-peptide-to-insulin ratio. n=10 control and n=9 GC-flattened mice. (b) Relationship between circulating C-peptide and insulin in the same mice. Linear regression equations are shown. (c–e) Representative liver immunoblots (c) and quantification of insulin-degrading enzyme (IDE) (d) and CEACAM1 (e) at Day 14. Protein abundance was normalised to vinculin. IDE abundance was reduced during GC-flattening, whereas CEACAM1 abundance was not significantly altered. n=8 mice per group. (f) Hepatic IDE activity in liver lysates. The time course shows fluorescence generated by IDE substrate cleavage, and the bar graph shows IDE activity calculated from the rate of fluorescence increase. n=4 mice per group. Data are mean ± SEM. Group comparisons in (a, d–f) were performed using unpaired two-tailed t tests with Welch’s correction for unequal variances. Asterisks indicate p values: *p<0.05, **p<0.01 and ***p<0.001. Only statistically significant pairwise comparisons are shown on the panels.

We therefore directly measured insulin clearance early during GC-flattening, when hyperinsulinaemia first develops. On Day 3, control and GC-flattened mice received the same intravenous dose of human insulin (0.125 U/kg), and circulating human insulin was measured using an assay that does not cross-react with endogenous mouse insulin (ESM Fig. 4). Following injection, circulating human insulin remained higher in GC-flattened mice than in controls, with approximately 2.4-fold greater insulin exposure between 2 and 10 min after injection (p=0.024; ESM Fig. 4), confirming slower insulin clearance. Thus, reduced insulin clearance is already present by Day 3, when the hyperinsulinaemic phenotype emerges.

We therefore examined whether this reduction in insulin clearance was accompanied by changes in hepatic proteins involved in insulin clearance, focusing on IDE and CEACAM1, which have both been implicated in insulin clearance [40–42]. Immunoblot analysis of liver lysates after 14 days of GC-flattening showed significantly reduced abundance of insulin-degrading enzyme (IDE) compared with controls (p = 0.038; Fig. 5c, d). In contrast, CEACAM1 abundance was not significantly altered (p = 0.245; Fig. 5c, e). Because reduced IDE abundance does not necessarily imply reduced enzymatic function, we directly measured IDE activity in liver extracts. GC-flattened liver extracts showed a slower increase in fluorescence in the IDE activity assay, and quantification confirmed significantly lower hepatic IDE activity in GC-flattened mice (Fig. 5f).

Together, these findings establish reduced insulin clearance as a second mechanism contributing to elevated circulating insulin during GC-flattening. Reduced hepatic IDE abundance and activity accompany this clearance defect, although these data do not establish that reduced IDE is its cause.

### Combined beta cell and hepatic GR deletion further reduces circulating insulin and worsens glycaemic control

To provide an independent genetic test of whether the elevated circulating insulin induced by GC-flattening is functionally required to maintain glucose homeostasis, we generated double-GRKO mice by administering hepatocyte-specific AAV8-TBG-iCre to beta cell GRKO mice (Fig. 6a). The MIP-CreERT and beta cell GRKO groups shown in Fig. 6 did not receive AAV. AAV8-TBG-LacZ-treated beta cell GRKO mice were analysed in a separate AAV-matched control cohort by ITT and GTT (ESM Fig. 5a–d). Because hepatic GR also affects glucose metabolism directly, we used the double-GRKO mice only to test the consequences of lowering circulating insulin further, not to determine why insulin clearance is reduced. Under control conditions, circulating insulin remained low in all three genotypes (Fig. 6b). During GC-flattening, circulating insulin was highest in MIP-CreERT controls, lower in beta cell GRKO mice and lowest in double-GRKO mice (Fig. 6c, d).

**Fig. 6.**
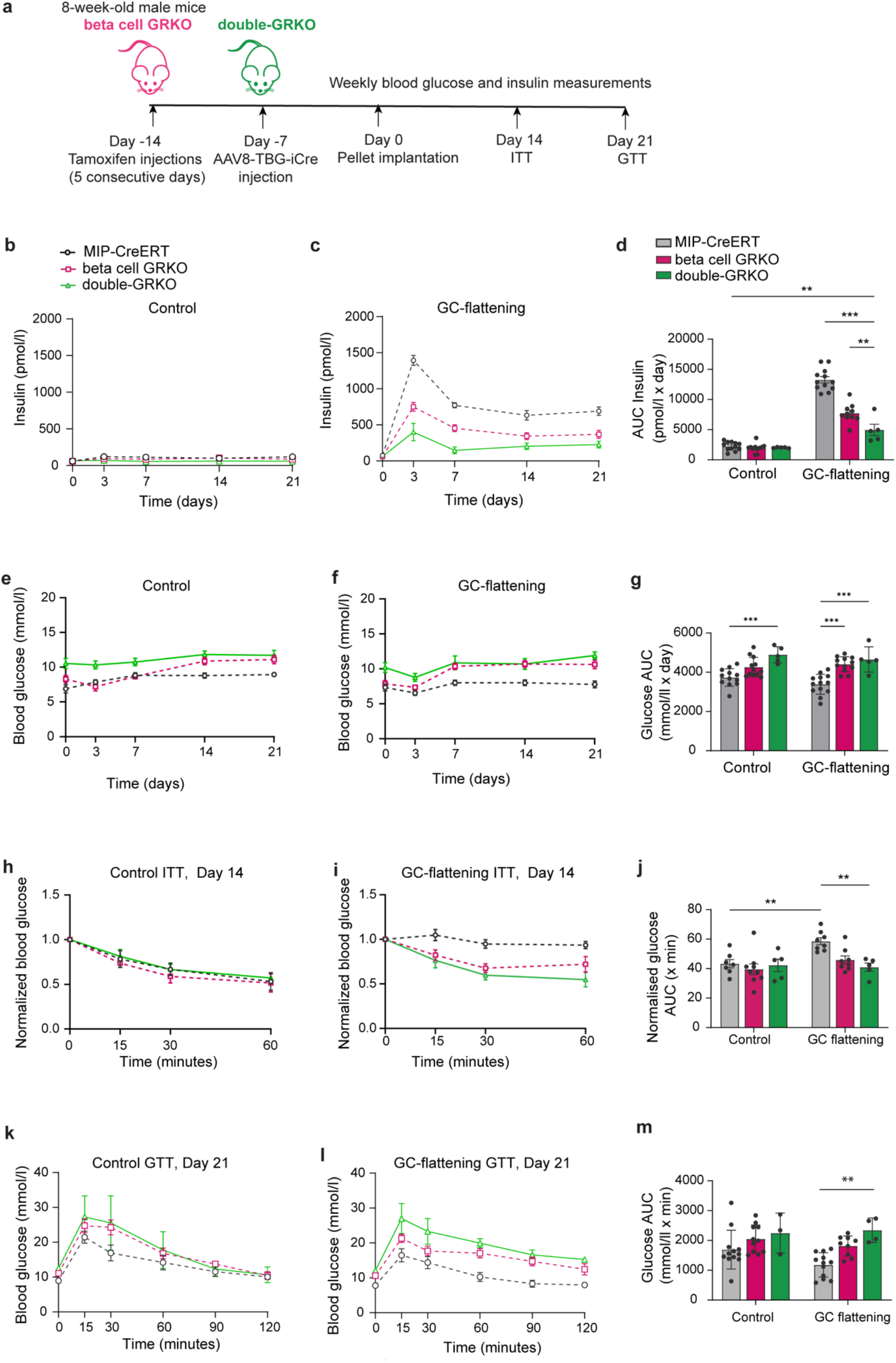
Combined beta cell and hepatic GR deletion further reduces circulating insulin and worsens glycaemic control during glucocorticoid rhythm disruption. (a) Experimental design. Adult beta cell GRKO mice were generated by tamoxifen treatment, and double-GRKO mice were generated by administering AAV8-TBG-iCre to beta cell GRKO mice 7 days before placebo or corticosterone pellet implantation. MIP-CreERT and beta cell GRKO mice did not receive AAV. AAV8-TBG-LacZ was administered to beta cell GRKO mice in separate control experiments (ESM Fig. 5). Hepatocyte GR deletion using the AAV8-TBG-iCre strategy was validated in the companion study [16]. The MIP-CreERT and beta cell GRKO groups are the same cohorts analysed in Figs 3 and 4. (b, c) Longitudinal fasting plasma insulin under control (b) and GC-flattened (c) conditions. (d) Insulin AUC over Days 0–21. During GC-flattening, insulin AUC was lower in beta cell GRKO than in MIP-CreERT mice and was reduced further in double-GRKO mice compared with both MIP-CreERT and beta cell GRKO mice. n=12 and 12 MIP-CreERT, n=11 and 10 beta cell GRKO, and n=5 and 5 double-GRKO mice under control and GC-flattened conditions, respectively. (e, f) Longitudinal fasting blood glucose under control (e) and GC-flattened (f) conditions. (g) Glucose AUC over Days 0–21. Under control conditions, glucose AUC was higher in double-GRKO mice than in MIP-CreERT controls. During GC-flattening, glucose AUC was higher in both beta cell GRKO and double-GRKO mice than in MIP-CreERT controls. n=12 and 13 MIP-CreERT, n=12 and 12 beta cell GRKO, and n=5 and 5 double-GRKO mice under control and GC-flattened conditions, respectively. (h, i) ITTs at Day 14 under control (h) and GC-flattened (i) conditions. Blood glucose is expressed relative to baseline. (j) ITT AUC. GC-flattening reduced insulin sensitivity in MIP-CreERT mice. Under GC-flattened conditions, double-GRKO mice were more insulin sensitive than MIP-CreERT controls, with beta cell GRKO mice showing an intermediate response. For ITTs, n=7, 9 and 5 for MIP-CreERT, beta cell GRKO and double-GRKO mice under control conditions and n=8, 8 and 5, respectively, under GC-flattened conditions. (k, l) GTTs at Day 21 under control (k) and GC-flattened (l) conditions. (m) Glucose AUC. During GC-flattening, glucose AUC was higher in double-GRKO mice than in MIP-CreERT controls. The difference between beta cell GRKO and MIP-CreERT mice did not reach statistical significance. Neither knockout genotype differed significantly from MIP-CreERT controls under control conditions. For GTTs, n=12, 11 and 3 for MIP-CreERT, beta cell GRKO and double-GRKO mice under control conditions and n=12, 10 and 4, respectively, under GC-flattened conditions. Data are mean ± SEM. Time-course data were analysed by two-way repeated-measures ANOVA with Tukey’s multiple-comparisons test. AUC measurements in (d, g, j) were analysed by two-way ANOVA with treatment and genotype as factors followed by Tukey’s multiple-comparisons test. GTT AUC in (m) was analysed by two-way ANOVA with treatment and genotype as factors followed by Šídák-adjusted planned comparisons. The treatment × genotype interaction was significant for insulin AUC (F(2,49)=40.0, p<0.0001) but not for glucose or ITT AUC. Asterisks indicate Tukey-or Šídák-adjusted multiple-comparison p values, as appropriate: *p<0.05, **p<0.01 and ***p<0.001. Only statistically significant pairwise comparisons are shown on the panels.

We examined how lower circulating insulin affected glucose homeostasis. MIP-CreERT controls maintained relatively stable fasting glucose during GC-flattening, whereas beta cell GRKO and double-GRKO mice developed higher glucose levels (Fig. 6e–g).

Importantly, the worsening of glycaemic control as circulating insulin was lowered was not caused by greater insulin resistance. During GC-flattening, MIP-CreERT controls were markedly insulin resistant, whereas beta cell GRKO and double-GRKO mice showed greater glucose lowering after insulin administration, with double-GRKO mice significantly more insulin sensitive than MIP-CreERT controls (Fig. 6h–j). This greater insulin sensitivity was also observed in an AAV-matched control cohort, in which double-GRKO mice were more insulin sensitive than beta cell GRKO mice receiving AAV8-TBG-LacZ (Šídák-adjusted p=0.011; ESM Fig. 5b), arguing against a nonspecific effect of AAV administration. Despite this improvement in insulin sensitivity, double-GRKO mice had worse glucose tolerance during GC-flattening (Fig. 6k–m).

Thus, further lowering circulating insulin by combined beta cell and hepatic GR deletion worsened glycaemic control despite improving insulin sensitivity. This is consistent with a model in which the GC-flattening-induced rise in insulin is matched to the accompanying insulin resistance and helps maintain glucose homeostasis.

## DISCUSSION

Our findings show that disruption of glucocorticoid timing can actively generate a hyperinsulinaemic state rather than hyperinsulinaemia arising solely as compensation for insulin resistance or increased glycaemia. During GC-flattening, beta cell GR-dependent enhancement of insulin secretion and reduced insulin clearance act together to increase circulating insulin.

Genetic attenuation of this insulin response improves insulin sensitivity but worsens glycaemic control, showing that the same hyperinsulinaemia that accompanies insulin resistance also becomes important for maintaining glucose homeostasis. Thus, GC-flattening reveals how an endocrine signal can drive hyperinsulinaemia and reshape the relationship between insulin secretion, insulin sensitivity and glycaemic control.

The beta cell experiments show that altered glucocorticoid timing produces a persistent change in insulin secretion. Islets from GC-flattened mice retained increased glucose-stimulated insulin secretion and Ca²⁺ responses after isolation, indicating that GC-flattening establishes a lasting change in beta cell glucose responsiveness rather than requiring the continued *in vivo* hormonal environment.

The increased beta cell glucose responsiveness during GC-flattening differs from the impaired glucose-stimulated insulin secretion often reported after sustained or pharmacological glucocorticoid exposure *in vitro* [20, 21], but is consistent with studies showing enhanced insulin secretion and glucose responsiveness after *in vivo* glucocorticoid treatment in rodents and humans [22–25, 43]. In those studies, the increased secretory response has generally been considered an adaptation to glucocorticoid-induced insulin resistance. Our findings show that glucocorticoids can also act directly on beta cells to enhance glucose-stimulated insulin secretion and that beta cell GR is required for much of this response during GC-flattening.

Importantly, our findings point to loss of the normal daily glucocorticoid trough as a key determinant of beta cell glucose responsiveness. GC-flattening raises this trough while maintaining mean glucocorticoid exposure near the physiological range, indicating that altered glucocorticoid timing can enhance beta cell responsiveness even without increased mean glucocorticoid exposure.

How loss of the daily glucocorticoid trough produces this persistent increase in beta cell glucose responsiveness through GR remains unclear. The response remains glucose dependent, arguing against nonspecific insulin hypersecretion and instead suggesting that GR alters the way beta cells respond to glucose. Potential mechanisms include changes in glucose sensing or metabolism, Ca²⁺ entry, or downstream amplification and exocytosis. One possibility, consistent with the proposal that elevation of the glucocorticoid trough increases daily GR occupancy [12], is that GC-flattening eliminates the normal interval of minimal GR activation and thereby produces more sustained GR-dependent transcription. Such sustained signalling could alter components of the glucose-sensing, Ca²⁺-handling or secretory machinery and thereby change the magnitude of glucose-stimulated insulin secretion [44]. More sustained GR signalling could also affect beta cell circadian organisation, because glucocorticoids can reset clocks in peripheral tissues [45] and the beta cell clock regulates insulin secretion [46]. Whether these transcriptional or circadian mechanisms account for the persistent secretory state will require direct testing.

The beta cell GR knockout experiments establish the physiological importance of this secretory response. Because GR was deleted in adult beta cells, the phenotype is unlikely to reflect developmental adaptation. During GC-flattening, beta cell GR deletion substantially reduced circulating insulin and worsened glucose tolerance despite similar or greater insulin sensitivity. This dissociation is important because it shows that the deterioration in glycaemic control is not caused by greater peripheral insulin resistance. Instead, the GR-dependent increase in insulin secretion is required to support glucose homeostasis during GC-flattening. The absence of a comparable secretory defect under control conditions further indicates that beta cell GR becomes particularly important when glucocorticoid rhythmicity is disrupted rather than being generally required for beta cell secretory capacity.

These experiments also show what happens when the GC-flattening-induced rise in insulin is reduced. Beta cell GR deletion lowers circulating insulin, improves insulin sensitivity and worsens glucose tolerance. Thus, the high insulin levels are not simply a marker of insulin resistance. They also help maintain glucose control once insulin resistance is present. At the same time, the improved insulin sensitivity after lowering insulin suggests that hyperinsulinaemia itself may contribute to the insulin-resistant state. Our data do not establish whether hyperinsulinaemia or insulin resistance develops first, because defects in muscle insulin signalling also appear rapidly [16]. Instead, they show that elevated insulin can both contribute to insulin resistance and help compensate for it.

Reduced insulin clearance also contributes to the rise in circulating insulin during GC-flattening. After the same intravenous dose of human insulin, circulating insulin remained higher in GC-flattened mice, confirming that insulin was removed from the circulation more slowly. This defect was already present by Day 3, when hyperinsulinaemia first appears. The reduced plasma C-peptide-to-insulin ratio at Day 14 provides additional evidence that reduced clearance persists during GC-flattening. GC-flattening also lowered hepatic IDE abundance and activity. However, this association does not establish that reduced IDE causes the clearance defect. Liver-specific deletion of *Ide* does not measurably reduce systemic insulin clearance in mice [47], arguing that reduced IDE alone is insufficient to explain the slower clearance. Together, these findings establish reduced insulin clearance as a second contributor to hyperinsulinaemia during GC-flattening and suggest that its molecular basis likely involves mechanisms beyond IDE.

The combined beta cell and hepatic GR deletion experiments provide an independent test of whether the high insulin levels are important for maintaining glycaemic control. Further lowering circulating insulin in double-GRKO mice worsened glycaemia despite greater insulin sensitivity. This result is especially informative because hepatic GR promotes gluconeogenic gene expression, so loss of hepatic GR would be expected to lower rather than raise blood glucose [48]. The worsening glycaemia therefore cannot be explained simply by a direct effect of hepatic GR deletion on glucose production. Together with the beta cell GRKO experiments, these findings strengthen the conclusion that the elevated insulin generated during GC-flattening is required to support glucose homeostasis. Because hepatic GR also regulates glucose metabolism directly, however, the double-GRKO experiments cannot determine how much of the hepatic contribution reflects altered insulin clearance.

Taken together, the secretion and clearance results suggest that GC-flattening raises circulating insulin through more than one mechanism and also induces insulin resistance. These processes may then reinforce one another. High insulin may contribute to insulin resistance, while insulin resistance increases the amount of insulin needed to maintain glycaemia. Our experiments do not establish which process occurs first or whether insulin resistance itself helps sustain the hyperinsulinaemia. Instead, they show that disrupted glucocorticoid timing can generate a state in which hyperinsulinaemia and insulin resistance coexist and become functionally linked. This model is summarised in Fig. 7.

**Fig. 7.**
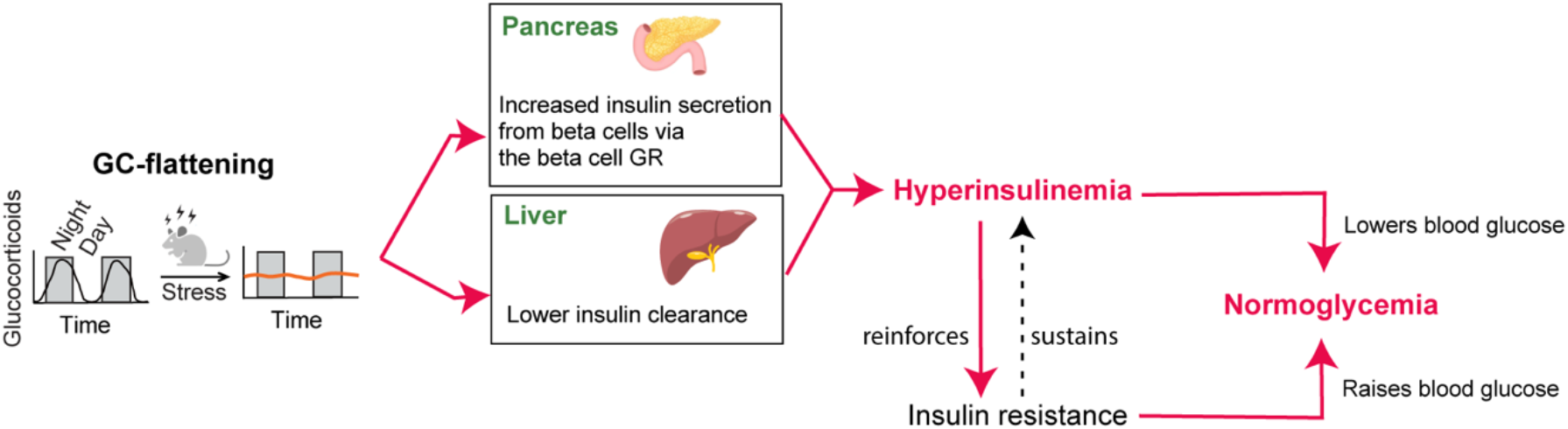
Proposed model of insulin regulation during glucocorticoid rhythm flattening. GC-flattening increases circulating insulin through beta cell GR-dependent enhancement of insulin secretion and reduced insulin clearance, accompanied by reduced hepatic IDE abundance and activity. GC-flattening also induces insulin resistance, particularly in skeletal muscle, which reduces insulin-mediated glucose lowering and may increase the amount of insulin needed to maintain glycaemia. Elevated insulin may in turn contribute to insulin resistance. The GC-flattening-induced rise in insulin is proposed to match the accompanying insulin resistance and thereby help maintain near-normal glycaemia. Solid arrows indicate relationships supported by the experimental data. The dashed arrow indicates the proposed contribution of insulin resistance to sustaining hyperinsulinaemia.

Several limitations remain. Although our data establish a role for beta cell GR in the increased secretory response during GC-flattening, the molecular targets linking GR signalling to altered beta cell stimulus-secretion coupling remain undefined. Direct insulin-clearance measurements were performed in a small cohort and did not resolve the initial distribution phase. The molecular basis of the clearance defect also remains unknown. Because hepatic GR directly regulates glucose metabolism, the double-GRKO experiments cannot determine how much of the hepatic contribution reflects altered insulin clearance. Finally, all experiments were performed in male mice, and whether the same mechanisms operate in females and humans remains to be determined.

More broadly, our findings show that hyperinsulinaemia need not arise solely as a compensatory response to insulin resistance or rising blood glucose. GC-flattening directly raises circulating insulin through increased beta cell secretion and reduced insulin clearance, while also inducing insulin resistance, particularly in skeletal muscle. Reducing the insulin response improves insulin sensitivity but worsens glycaemic control, revealing two seemingly opposing roles for the elevated insulin: it contributes to the insulin-resistant state, yet once that resistance is present, it is also required to maintain glucose homeostasis. By demonstrating both roles within the same physiological state, our findings explain how apparently conflicting models in which hyperinsulinaemia is viewed as either a cause or a consequence of insulin resistance can both be valid.

## ACKNOWLEDGEMENTS

We thank T. Meyer (Weill Cornell Medicine, New York, NY, USA), A. Tengholm (Uppsala University, Uppsala, Sweden) and M. Merrins (Yale University, New Haven, CT, USA) for valuable scientific discussions and advice throughout the study. We thank A. Tengholm, M. Merrins, D. Scott and L. Katz (Icahn School of Medicine at Mount Sinai, New York, NY, USA) for critical reading of the manuscript. We also thank members of the Teruel and Meyer laboratories for helpful discussions, and S. Chen, A. C. N. Chong, L. Yammine and H. Foster for helpful advice.

## AI-assisted technology disclosure

ChatGPT (OpenAI) was used during manuscript preparation to assist with editing, organisation and refinement of the text and to improve clarity of presentation. All scientific content, data analyses, interpretations, references and final wording were independently reviewed and verified by the authors. AI was not used to generate experimental data or figures.

## AUTHORS’ RELATIONSHIPS AND ACTIVITIES

The authors declare that there are no relationships or activities that might bias, or be perceived to bias, their work.

## CONTRIBUTION STATEMENT

JW and MNT designed the study. JW, ASA, SS, AA and JL acquired, analysed and interpreted the data. MNT analysed and interpreted the data, provided supervision and acquired funding. JW and MNT drafted the manuscript, and all authors revised it critically for important intellectual content and approved the final version. MNT is the guarantor of this work and, as such, had full access to all the data in the study and takes responsibility for the integrity of the data and the accuracy of the data analysis

## DATA AVAILABILITY

The datasets supporting the findings of this study will be available from the corresponding author, Mary N. Teruel, from the date of publication for at least 5 years. Data will be shared electronically on reasonable written request with researchers for non-commercial research purposes; requests should identify the data sought and the planned analyses. No additional restrictions apply. No custom code was generated in this study.

## ABBREVIATIONS

AAV: Adeno-associated virus
CEACAM1: Carcinoembryonic antigen-related cell adhesion molecule 1
GC: Glucocorticoid
GR: Glucocorticoid receptor
GRKO: Glucocorticoid receptor knockout
GSIS: Glucose-stimulated insulin secretion
GTT: Glucose tolerance test
HFD: High-fat diet
IDE: Insulin-degrading enzyme
ITT: Insulin tolerance test
MACS: Mild autonomous cortisol secretion

