## Supplementary Material for "Glucocorticoid rhythm disruption drives hyperinsulinaemia in mice through beta cell glucocorticoid receptor signalling"

### Electronic Supplementary Material (ESM)

*Detailed protocols for the methods summarised in the main-text Methods section, together with the supplementary figure legends. Mouse strain, reagent and antibody details are given here in full.*

#### Experimental model

**Ethics.** All animal procedures were approved by the Institutional Animal Care and Use Committee (IACUC) of Weill Cornell Medical College (protocol number 2020-0001) and were performed in accordance with the NIH Guide for the Care and Use of Laboratory Animals and applicable institutional and federal regulations.

**Animals.** Male mice aged 8–14 weeks at the time of pellet implantation were used for all experiments. Wild-type animals were C57BL/6J mice, and the genetically modified lines (beta cell GRKO, double-GRKO and their littermate controls) are described under “Mouse strains and generation…” and “Generation of beta cell and liver-specific GR double knockout mice”, where provenance and genotyping are given. Mice were maintained under specific-pathogen-free conditions and were experimentally naïve before the procedures described below, other than tamoxifen and/or AAV administration where applicable.

Housing and husbandry. Animals were housed in the Weill Cornell Medicine animal facility in individually ventilated cages at up to five mice per cage under a 12-h light/dark cycle (lights on at 06:00), at 22 ± 2 °C and 30–70% relative humidity. Water and standard chow (PicoLab Rodent Diet 20, 5053, Lab Diets, St. Louis, MO, USA) were available ad libitum except during designated fasting periods. Cages were provided with corncob bedding and nesting material. Mice were acclimatised to the facility for 7 days before any procedure. For diet-induced obesity studies, mice were fed a high-fat diet containing 60% kcal from fat (D12492, Research Diets, New Brunswick, NJ, USA) ad libitum.

Randomisation and blinding. Animals were individually identified by ear tags. Before pellet implantation, experimental and control groups were matched for baseline body weight using block randomisation to ensure equivalent starting weights. To minimise potential confounders, experimental and control animals were co-housed whenever possible, cage position on the rack was not assigned by group, and measurements and dissections were performed alternately between corticosterone-treated and control animals. Investigators were blinded to treatment and genotype during outcome assessment and data analysis. Samples and image or data files were coded and were unblinded only after quantification was complete. Blinding was not possible during pellet implantation and tamoxifen or AAV administration because these procedures require knowledge of the agent being delivered.

**Inclusion and exclusion criteria.** Inclusion and exclusion criteria were defined prospectively. Animals were eligible if they met the strain, sex and age criteria above and underwent successful pellet implantation and, where applicable, successful tamoxifen-induced and/or AAV-mediated recombination. Pre-specified exclusion criteria were loss or extrusion of the pellet, surgical complications, or illness unrelated to treatment. No data points were excluded from any analysis. The exact number of animals or biological replicates analysed in each group is reported in the corresponding figure legend.

Animal care, monitoring and humane endpoints. Mice were monitored at least once daily by animal-facility and research staff, with additional monitoring after surgical pellet implantation.

Pellet implantation was performed under isoflurane anaesthesia as described under "Corticosterone administration in mice". Immediately after induction of anaesthesia, mice received perioperative analgesia in accordance with the approved IACUC protocol, together with local infiltration of bupivacaine (Marcaine 0.25%, 0.1 ml) at the incision site. Additional analgesia was provided if signs of pain were observed, including altered ambulation, decreased grooming or rearing, decreased socialisation, a dirty coat or lethargy. Body weight and general condition, including posture, coat, activity and hydration, were assessed weekly. Pellet-implanted mice were additionally observed daily during the first week after implantation to monitor wound healing and every second day thereafter. Pre-defined humane endpoints were weight loss exceeding 10% of baseline body weight, lethargy, dehydration, signs of pain or distress, or failure to thrive. No animal reached a humane endpoint during the study, and no unexpected adverse events were observed. At the study endpoint, mice were euthanised by $CO_2$ asphyxiation followed by cervical dislocation, as approved by the IACUC.

#### Islet isolation and culture

Pancreatic islets were isolated by collagenase digestion. The pancreas was perfused through the common bile duct with HBSS containing 1.7 mg/ml collagenase P (Sigma-Aldrich), excised and digested for 21 min at 37 °C with intermittent gentle agitation. Digestion was stopped using ice-cold HBSS containing 10% FBS. The tissue was filtered through a mesh strainer and purified by Histopaque density-gradient centrifugation. Islets were handpicked under a stereomicroscope and recovered overnight at 37 °C in a humidified 5% $CO_2$ incubator under the assay-specific culture conditions described below.

#### Mouse strains and generation of beta cell–specific glucocorticoid receptor knockout mice (beta cell GRKO)

C57BL/6J mice (stock #000664), Nr3c1fl/fl mice (hereafter GRfl/fl, stock #021021) and MIP-CreERT mice (stock #024709) were obtained from The Jackson Laboratory. Beta cell–specific glucocorticoid receptor knockout mice (beta cell GRKO) were generated using a two-generation breeding strategy. Female GRfl/fl mice were crossed with male MIP-CreERT mice to generate MIP-CreERT;GRfl/+ offspring, which were subsequently bred with GRfl/fl mice to produce MIP-CreERT;GRfl/fl mice, hereafter referred to as beta cell GRKO. Male littermate GRfl/fl mice lacking the Cre transgene and MIP-CreERT mice were used as controls.

All strains were maintained on a C57BL/6J genetic background. GRfl/fl control mice were derived from the same two-generation breeding strategy described above. MIP-CreERT mice were maintained by crossing male carriers with wild-type C57BL/6J females. Pups were genotyped by PCR (Transnetyx, Inc., Cordova, TN) within four weeks of birth.

#### Tamoxifen administration

Cre-mediated recombination in beta cells was induced by tamoxifen. Tamoxifen (Sigma-Aldrich, St. Louis, MO, USA) was dissolved in sterile corn oil (15 mg/ml) and incubated overnight at 37 °C in a light-protected container. Two dosing schedules were used, both delivering tamoxifen intraperitoneally at 75 mg/kg body weight to adult mice aged 8–10 weeks. For beta cell GRKO and double-GRKO phenotyping experiments, including the flow-cytometric validation in ESM Fig. 3C, mice received one injection daily for five consecutive days (days 1–5). A reduced schedule of three injections on alternate days (days 1, 3 and 5) was used only for the immunofluorescence validation in ESM Fig. 3A, B to test whether fewer doses were sufficient for efficient recombination. Each schedule was followed by a 7-day washout period to allow for drug clearance and protein turnover. Control animals received identical injections under the corresponding schedule. Phenotypic analyses were performed after the washout period, and GR-deletion efficiency was validated 3–5 weeks after the final injection.

#### Generation of beta cell and liver-specific GR double knockout mice (double-GRKO)

Beta cell and hepatocyte glucocorticoid receptor double-knockout mice (double-GRKO) were generated using a sequential strategy. GR deletion in beta cells was first induced by administering tamoxifen to beta cell GRKO mice as described above. Double-GRKO mice were then generated in a separate cohort by retro-orbital injection of AAV8-TBG-iCre (Vector Biolabs, VB1724), with 100 µl containing 2 × $10^{11}$ genome copies diluted in sterile PBS, 7 days after starting the tamoxifen injections. Pellet implantation and phenotypic experiments were started 7 days after AAV8-TBG-iCre injection. The MIP-CreERT and beta cell GRKO groups shown in Fig. 6 were the same cohorts used in Figs 3 and 4 and did not receive AAV. AAV8-TBG-LacZ (Vector Biolabs, 70003-S) was used in separate AAV control experiments, including the direct insulin-clearance experiment shown in ESM Fig. 4, but was not used as the comparator treatment for the Fig. 6 MIP-CreERT or beta cell GRKO groups. Hepatocyte GR deletion using the AAV8-TBG-iCre strategy was validated in the companion GC-flattening manuscript [1].

#### Ins1-Cre;GCaMP6f mice (beta cell $Ca^{2+}$ reporter)

To generate mice expressing the calcium reporter GCaMP6f specifically in pancreatic beta cells, homozygous Ins1-Cre mice (JAX #026801) were crossed with homozygous GCaMP6f reporter mice (JAX #028865). In the resulting offspring, Cre recombinase expressed under the insulin promoter drives beta cell–specific expression of GCaMP6f via excision of a loxP-flanked stop cassette.

#### Fasting conditions and sample collection

Throughout this manuscript, “fasted” mice refer to animals placed in clean cages with no access to food, food dust or faeces in the bedding for 5 h during the light cycle, typically 09:00–14:00. Water was freely available during fasting. Unless otherwise noted, mice were fasted before blood and tissue collection. Blood was collected by tail snip or cardiac puncture into EDTA-coated tubes, centrifuged at 6,000 × g for 20 min at 4 °C, and plasma was stored at −20 °C. Dissections were performed between 14:00 and 18:00, alternating between corticosterone-treated and control mice. Tissues were either fixed in buffered zinc formalin (Cancer Diagnostics, 171) for paraffin embedding or snap-frozen in liquid nitrogen and stored at −80 °C.

#### Fasting glucose and insulin measurements

For fasting glucose and insulin measurements, mice were fasted for 5 h as described above, with sample collection performed between 14:00 and 18:00. Blood was drawn by tail snip, and glucose was measured using a glucometer (Diathrive, Salt Lake City, UT, USA). An additional 30 µl of blood was collected into EDTA-coated tubes for plasma insulin measurements. For subsequent weekly blood collections, samples were obtained by removing the scab from the previous draw site. Plasma insulin was measured using the Ultra-Sensitive Mouse Insulin ELISA Kit (90080, Crystal Chem, Elk Grove Village, IL, USA) according to the manufacturer’s instructions.

#### Tolerance tests

Before glucose tolerance tests (GTTs) and insulin tolerance tests (ITTs), mice were fasted for 5 h starting at 09:00 as described under “Fasting conditions and sample collection”. For GTTs, mice received glucose (2 g/kg body weight, Sigma-Aldrich G8270) by intraperitoneal injection.

For ITTs, mice received human insulin (0.5 U/kg body weight, Humulin R U-100, Eli Lilly) by intraperitoneal injection. Blood glucose was measured immediately before injection and at 15, 30 and 60 min after insulin for ITTs, and at 15, 30, 60, 90 and 120 min after glucose for GTTs.

**Corticosterone administration in mice**

Mice were implanted subcutaneously with corticosterone pellets designed for either 21-day or 60-day release. The 21-day pellet contained 5 mg corticosterone (G-111), and the 60-day formulation contained 15 mg total corticosterone (SG-111, Innovative Research of America, Sarasota, FL, USA). Placebo pellets (C-111 or SC-111) served as controls. Mice were anaesthetised by continuous inhalation of isoflurane. Once anaesthetised, a small incision was made on the lateral side of the neck and the pellet was inserted approximately 2 cm from the incision using forceps or a trocar. Mice weighed 24.1 ± 1.2 g on average, corresponding to a corticosterone dose of approximately 10 mg $kg^{-1}$ $day^{-1}$.

**Dynamic glucose-stimulated insulin secretion (GSIS) assay on isolated islets**

Dynamic glucose-stimulated insulin secretion was measured in three experimental settings. First, islets isolated from wild-type C57BL/6J mice were recovered overnight and treated for 24 h with corticosterone (280 nmol/l) or DMSO vehicle. Second, islets were isolated from wild-type mice 2 days after corticosterone or placebo pellet implantation. Third, islets were isolated from MIP-CreERT control and beta cell GRKO mice 2 days after corticosterone or placebo pellet implantation. Islets isolated after in vivo treatment were recovered overnight before perifusion. A PERI-4.02 perifusion system (Biorep Technologies, Miami, FL, USA) was primed with Krebs buffers containing 3, 8 or 20 mmol/l glucose. Islets were preincubated for 2 h in Krebs buffer containing 3 mmol/l glucose and loaded into individual columns at 100 islet equivalents (IEQ) per column. Islets were perifused sequentially with 3 mmol/l glucose for 40 min, 8 mmol/l glucose for 20 min, 3 mmol/l glucose for 20 min and 20 mmol/l glucose for 20 min. Buffer was delivered at 80 µl/min, and perifusate was collected every 2 min into 96-well plates. Perifusate samples were stored at −20 °C until insulin was quantified using the Lumit Insulin Immunoassay (W8012, Promega, Madison, WI, USA).

**Islet insulin content after glucose or KCl exposure**

To determine whether enhanced secretion during GC-flattening was accompanied by depletion of islet insulin stores, islets were isolated from placebo- and corticosterone-pelleted mice on Day 14 and recovered overnight at 37 °C in 5% $CO_2$. The assay was adapted from Yammine et al. [2] and Slepchenko et al. [3]. For each biological replicate, islets from three mice were pooled. Forty islet equivalents (IEQ) were analysed per condition, with 1 IEQ defined as an islet with a diameter of 150 µm. Islets were preincubated for 2 h at 37 °C in Krebs buffer containing 20 mmol/l HEPES, 1 mmol/l $MgSO_4$, 119 mmol/l NaCl, 4.6 mmol/l KCl, 0.4 mmol/l $KH_2PO_4$, 2 mmol/l $CaCl_2$, 5 mmol/l $NaHCO_3$, 0.05% BSA and 2.7 mmol/l glucose, adjusted to pH 7.4. Islets were then incubated for 45 min at 37 °C in the same buffer containing either 2.7 mmol/l glucose, 16.7 mmol/l glucose or 30 mmol/l KCl together with 2.7 mmol/l glucose.

Immediately after incubation, islets were collected in ice-cold extraction buffer containing 0.18 mol/l HCl in 70% ethanol. Samples were vortexed, incubated on ice for 30 min, vortexed again and maintained at 4 °C for 24 h. Samples were then centrifuged, and the supernatants were collected and stored at −20 °C. Insulin was quantified using the Ultra-Sensitive Mouse Insulin ELISA Kit (90080, Crystal Chem) according to the manufacturer’s instructions. Islet insulin content was expressed as the insulin concentration in the extract per 40 IEQ. Three independent biological replicates were analysed per treatment group, with each replicate containing pooled islets from three mice.

#### C-peptide:insulin ratio

For measurements after 14 days of treatment, plasma was collected following the 5-h fast. Insulin was measured by radioimmunoassay and C-peptide using a Luminex assay. The C-peptide:insulin ratio was calculated from molar concentrations.

#### Direct insulin clearance

Direct insulin clearance was measured on Day 3 of treatment in GRfl/fl mice that had received AAV8-TBG-LacZ. Control and GC-flattened mice received human insulin (0.125 U/kg body weight) by injection into the lateral tail vein. Plasma was collected immediately before injection and at 2, 5, 10 and 15 min after injection. Human insulin was quantified using a chemiluminescent immunoassay that detects human insulin without cross-reactivity to endogenous mouse insulin. Insulin exposure was quantified as the area under the plasma human-insulin concentration curve from 2 to 10 min using the linear trapezoidal rule, the widest interval sampled in every animal. Because all mice received the same dose per unit body weight, relative clearance was calculated for each mouse as the geometric mean 2–10 min AUC of the control group divided by that mouse's 2–10 min AUC. Absolute metabolic clearance rate was not calculated because the first post-injection sample was collected at 2 min and the initial distribution phase was therefore unresolved.

#### Calcium imaging

Calcium imaging was performed using intact islets from Ins1-Cre;GCaMP6f mice generated as described above.

For direct corticosterone-treatment experiments, islets were isolated from male mice aged 8–12 weeks and recovered overnight at 37 °C in 5% $CO_2$ in RPMI-1640 containing 11 mmol/l glucose, 2 mmol/l L-glutamine, 100 IU/ml penicillin, 100 μg/ml streptomycin and 10% fetal bovine serum. Islets were then treated for 24 h in the same medium containing 5 mmol/l glucose and either corticosterone (280 nmol/l) or DMSO vehicle.

For in vivo GC-flattening experiments, mice were implanted with corticosterone or placebo pellets. Islets were isolated 2 days later and recovered overnight in medium containing 11 mmol/l glucose. Islets from one or two mice were pooled for each biological replicate.

Before imaging, islets were incubated for 2 h in Krebs-HEPES-bicarbonate buffer containing 130 mmol/l NaCl, 3.6 mmol/l KCl, 1.5 mmol/l $CaCl_2$, 0.5 mmol/l $MgSO_4$, 0.5 mmol/l $NaH_2PO_4$, 24 mmol/l $NaHCO_3$, 10 mmol/l HEPES and 3 mmol/l glucose, adjusted to pH 7.4. Islets were transferred to an imaging chamber containing the same buffer and allowed to settle for 10 min. They were then perifused sequentially with 3 mmol/l glucose for 9 min, 8 mmol/l glucose for 20 min, 3 mmol/l glucose for 20 min and 20 mmol/l glucose for 16 min. Imaging was performed at 37 °C using a 3i Yokogawa spinning-disk confocal microscope equipped with a 20×/0.8 Plan Achromat objective. GCaMP6f was excited using the 488-nm laser line. Images were acquired every 500 ms using a 400-ms exposure followed by a 100-ms interval. Images were analysed in ImageJ. A region of interest was drawn around each intact islet, and whole-islet mean fluorescence intensity was measured over time. Each trace was normalised to its first time point. Averaged traces contained 5–15 individual islet traces. Exact numbers of biological replicates and islets analysed are provided in the corresponding figure legends.

#### Insulin-degrading enzyme activity assay

Hepatic insulin-degrading enzyme (IDE) activity was measured using the SensoLyte 520 IDE Activity Assay Kit (AS72231, AnaSpec, Fremont, CA, USA). Frozen liver tissue (25 mg) was homogenised in assay buffer on ice. Homogenates were incubated on ice for 15 min and centrifuged at 10,000 × g for 15 min at 4 °C. Supernatants were collected and stored at −80 °C until analysis. Lysates were incubated with the FRET-based IDE substrate according to the manufacturer's instructions. Substrate cleavage releases 5-FAM fluorescence, which was measured at excitation/emission wavelengths of 490/520 nm every 5 min for 60 min at 37 °C using a microplate reader. IDE activity was expressed as relative fluorescence intensity over time.

#### Immunoblotting

Liver tissue was homogenised in ice-cold lysis buffer containing protease and phosphatase inhibitors, and lysates were cleared by centrifugation. Protein concentration was measured by BCA assay. Equal amounts of protein (35 µg per lane) were separated by SDS-PAGE on 10% gels and transferred to PVDF membranes. Membranes were blocked and incubated overnight at 4 °C with antibodies against IDE, CEACAM1 or vinculin, followed by the appropriate fluorescent secondary antibodies. Immunoreactive bands were visualised and quantified using LI-COR Image Studio Lite. IDE and CEACAM1 signals were normalised to vinculin. Uncropped immunoblot images are provided in a separate source-image file. Primary antibodies were IDE (Abcam, ab32216, RRID: AB_775686), CEACAM1 (Cell Signaling Technology, 14771S, RRID: AB_2798605) and vinculin (Thermo Fisher Scientific, 14-9777-80, RRID: AB_2573027), each used at 1:1,000. Goat anti-rabbit Alexa Fluor 680 secondary antibody (Thermo Fisher Scientific, A21109, RRID: AB_2535758) was used at 1:5,000.

#### Whole-mount islet immunofluorescence to validate beta cell GR deletion

Islets isolated from tamoxifen-treated GRfl/fl control mice and beta cell GRKO mice were cultured for 72 h, collected in BSA-coated tubes and fixed in 4% paraformaldehyde (Electron Microscopy Sciences, 15710-S) for 2 h at room temperature with rotation. Fixed islets were washed twice in PBS, transferred to low-attachment U-bottom 96-well plates at 5–10 islets per well and permeabilised in PBS containing 0.3% Triton X-100 for 1 h. Islets were blocked for 1 h in PBS containing 1% BSA, 10% FBS and 0.1% Triton X-100. Islets were incubated overnight at 4 °C with antibodies against insulin (guinea pig, Agilent/Dako, A0564, 1:5) and GR (rabbit monoclonal clone D6H2L, Cell Signaling Technology, 12041, 1:250). After washing in PBS containing 0.1% Tween-20, islets were incubated for 2 h at room temperature with Alexa Fluor 488 goat anti-guinea pig IgG (Invitrogen, A-11073, 1:1,000), Alexa Fluor 647 goat anti-rabbit IgG (Invitrogen, A-21245, 1:500) and Hoechst 33342 (Invitrogen, 62249, 1:10,000). Islets were mounted in Fluoromount-G (Electron Microscopy Sciences, 17984-25) and imaged by confocal microscopy using a 20× objective without binning. Beta cell GR-deletion efficiency was quantified in Fiji (ImageJ, NIH) from 18 islets from one beta cell GRKO mouse and eight islets from one GRfl/fl control mouse. Insulin-positive cells were identified as beta cells and Hoechst staining was used to identify nuclei. Within the insulin-positive population, cells were manually scored as GR-positive or GR-negative based on detectable nuclear GR immunofluorescence. Deletion efficiency was calculated for each islet as the percentage of insulin-positive cells lacking detectable nuclear GR signal.

#### Flow cytometry to validate beta cell GR deletion

Islets were isolated 3–5 weeks after tamoxifen administration and dispersed into single-cell suspensions using 0.05% trypsin-EDTA at 37 °C with periodic trituration. Cells were washed in

PBS containing 3% BSA and transferred to round-bottom 96-well plates. Viability was assessed using Live/Dead Blue, and macrophages were labelled with F4/80-PE (BioLegend, 123110, 1:500). Cells were fixed using BD Phosflow Lyse/Fix Buffer and permeabilised using BD Phosflow Perm Buffer III. Cells were incubated for 1 h with antibodies against insulin (DAKO, A0564, 1:50) and GR (Cell Signaling Technology, 12041, 1:500), followed by fluorophore-conjugated secondary antibodies, including goat anti-rabbit Alexa Fluor 647 (Invitrogen, A-21245, 1:500). Compensation controls were prepared using UltraComp eBeads stained individually with each fluorophore-conjugated antibody. Beta cells were identified as live, single, F4/80-negative, insulin-positive events. The insulin-positive gate was established using a fluorescence-minus-one control stained with secondary antibody alone. Within the beta cell population, the boundary between GR-positive and GR-negative cells was defined using a GR fluorescence-minus-one control and confirmed using islet cells from tamoxifen-treated GRfl/fl mice. Deletion efficiency was expressed as the percentage of GR-negative cells within the insulin-positive population. Median GR fluorescence intensity was also recorded as a gating-independent measure of receptor loss. Flow-cytometric validation was performed in one mouse per genotype for GRfl/fl, MIP-CreERT and beta cell GRKO mice. RRIDs were AB_893486 for F4/80-PE, AB_10013624 for insulin, AB_2631286 for GR and AB_2535813 for goat anti-rabbit Alexa Fluor 647.

#### Statistical analysis

Data are presented as mean ± SEM. Group comparisons used one- or two-way ANOVA with Tukey's or Šídák's multiple-comparisons tests and unpaired two-tailed Student's t tests for two-group or area-under-the-curve comparisons, as specified in the corresponding figure legends. A value of $p < 0.05$ was considered statistically significant. Analyses were performed in GraphPad Prism v11 (GraphPad Software, San Diego, CA, USA).

**ESM figures**

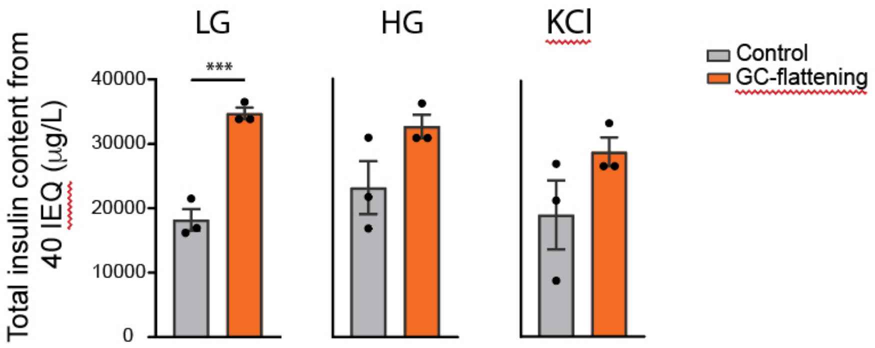


**ESM Fig. 1. GC-flattening does not deplete islet insulin stores after glucose or KCl exposure.**

Insulin content remaining in islets isolated from control and GC-flattened mice on Day 14 was measured by acid–ethanol extraction after incubation under low-glucose conditions (LG, 2.7 mmol/l glucose), high-glucose conditions (HG, 16.7 mmol/l glucose) or KCl-stimulated conditions (KCl, 30 mmol/l KCl with 2.7 mmol/l glucose). Insulin content was not reduced in islets from GC-flattened mice under any condition and was significantly increased after low-glucose exposure. These findings indicate that enhanced insulin secretion during GC-flattening is not attributable to depletion of islet insulin stores. Data are mean ± SEM. Each point represents one independent biological replicate (n = 3 per group), comprising islets pooled from three mice, with 40 IEQ analysed per condition. Groups were compared within each condition using unpaired two-tailed Student's t tests. Asterisks indicate p values: ***$p<0.001$. Exact p values were LG, $p=0.0009$, HG, $p=0.1023$ and KCl, $p=0.1666$.

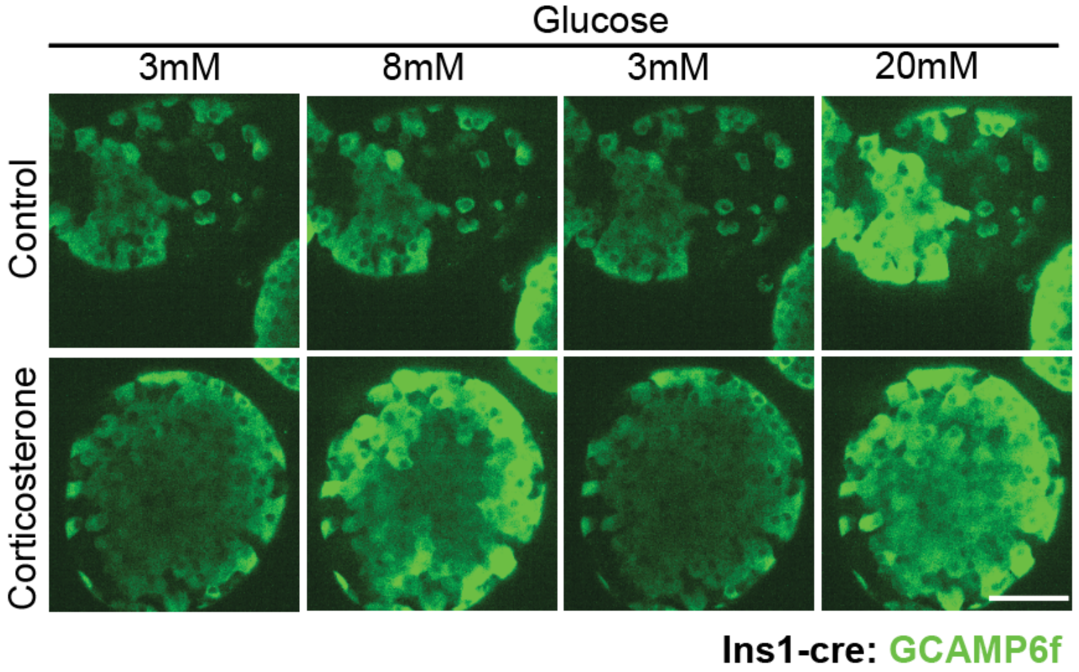


**ESM Fig. 2. Corticosterone increases glucose-stimulated beta cell $Ca^{2+}$ responses.**
Representative GCaMP6f fluorescence images of islets from Ins1-Cre;GCaMP6f mice at the indicated glucose concentrations, comparing DMSO vehicle and corticosterone-treated (280 nmol/l, 24 h) conditions. Corticosterone-treated islets show increased GCaMP6f fluorescence under stimulatory glucose concentrations. Scale bar, 50 μm.

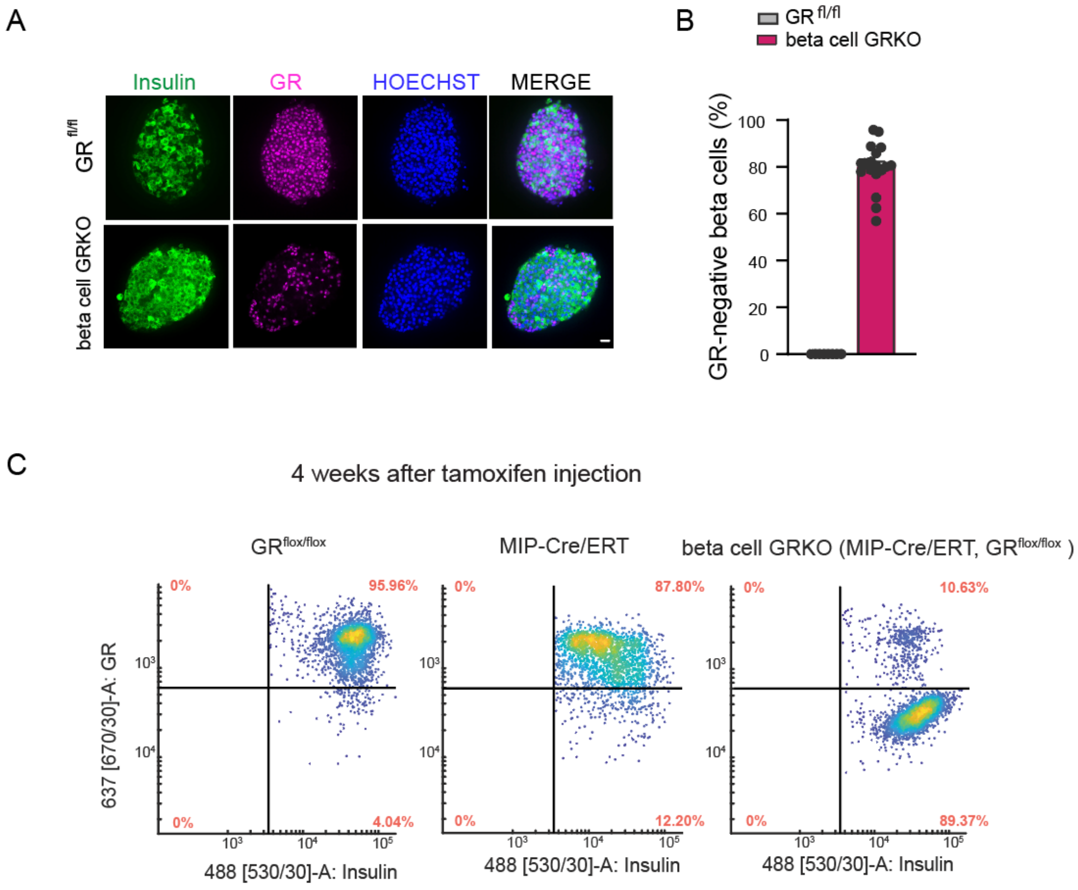


**ESM Fig. 3. Validation of beta cell–specific glucocorticoid receptor deletion.**

(A) Representative whole-mount immunofluorescence images of isolated islets from GRfl/fl control and beta cell GRKO mice stained for insulin (green), glucocorticoid receptor (GR, magenta) and nuclei (Hoechst, blue), with the merged image shown at right (scale bar, 20 μm). Both genotypes received the reduced tamoxifen schedule of three injections on alternate days (days 1, 3 and 5). GRfl/fl control mice lack the MIP-CreERT transgene, whereas beta cell GRKO islets show markedly reduced GR immunoreactivity. (B) Beta cell GR-deletion efficiency quantified from the immunofluorescence images in (A), corresponding to approximately 80% loss of beta cell GR. Points represent individual islets (18 islets from one beta cell GRKO mouse and eight islets from one GRfl/fl control mouse), and bars show mean ± SEM. No inferential statistical test was performed because biological n = 1 per genotype. (C) Flow cytometry of GR and insulin co-staining in dispersed islet cells from GRfl/fl, MIP-CreERT and beta cell GRKO mice after the standard five-consecutive-day tamoxifen schedule, analysed four weeks after the final injection (n = 1 mouse per genotype). Cells were gated on insulin-positive beta cells, and the GR-positive/GR-negative boundary was defined using a fluorescence-minus-one control. In the beta cell GRKO mouse, 89% of insulin-positive cells were GR-negative, compared with 4% in the GRfl/fl mouse and 12% in the MIP-CreERT mouse. Together, immunofluorescence and flow cytometry confirmed efficient beta cell GR deletion.

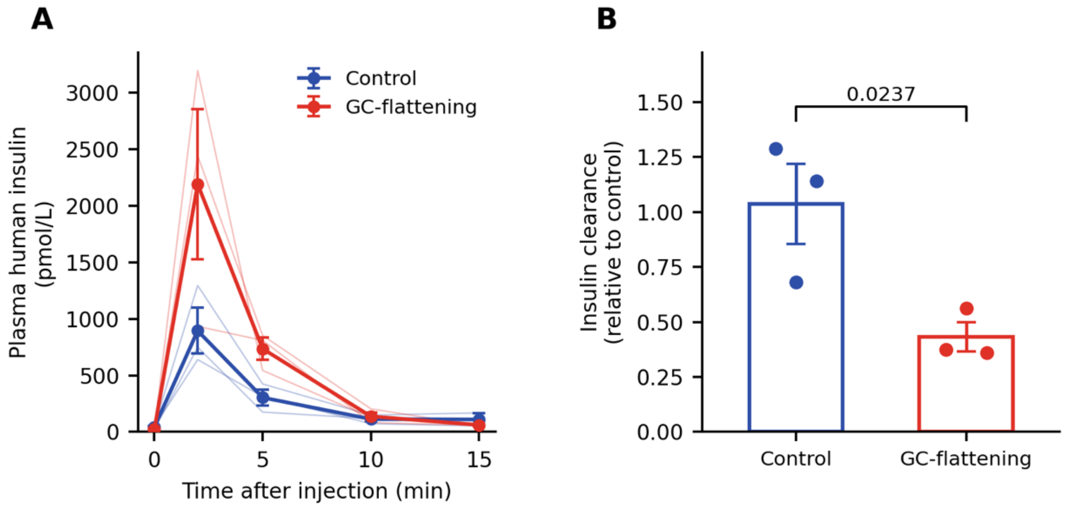


**ESM Fig. 4. Direct measurement of insulin clearance on Day 3 of GC-flattening.**

(A) Plasma human insulin concentration after a single injection of human insulin (0.125 U/kg body weight) into the lateral tail vein on Day 3 of treatment. Human insulin was measured using a chemiluminescent immunoassay that does not cross-react with mouse insulin, so endogenous mouse insulin does not contribute to the measured signal. Thin lines show individual mice and bold symbols show the group mean. n = 3 mice per group. The 15-min time point represents two control mice because one sample was not available. (B) Insulin clearance relative to control. Because all mice received the same dose per unit body weight, relative clearance was calculated as the geometric mean 2–10 min AUC of the control group divided by the 2–10 min AUC of each mouse. AUC was integrated using the linear trapezoidal rule. Insulin clearance in GC-flattened mice was 0.42 of control, corresponding to a 2.4-fold higher human-insulin AUC. Absolute metabolic clearance rate is not reported because the first post-injection sample was collected at 2 min and the initial distribution phase was unresolved. Mice in this experiment were GRfl/fl animals that received AAV8-TBG-LacZ. Data are mean ± SEM. Relative clearance was compared using an unpaired two-tailed Student’s t test. The exact p value is shown.

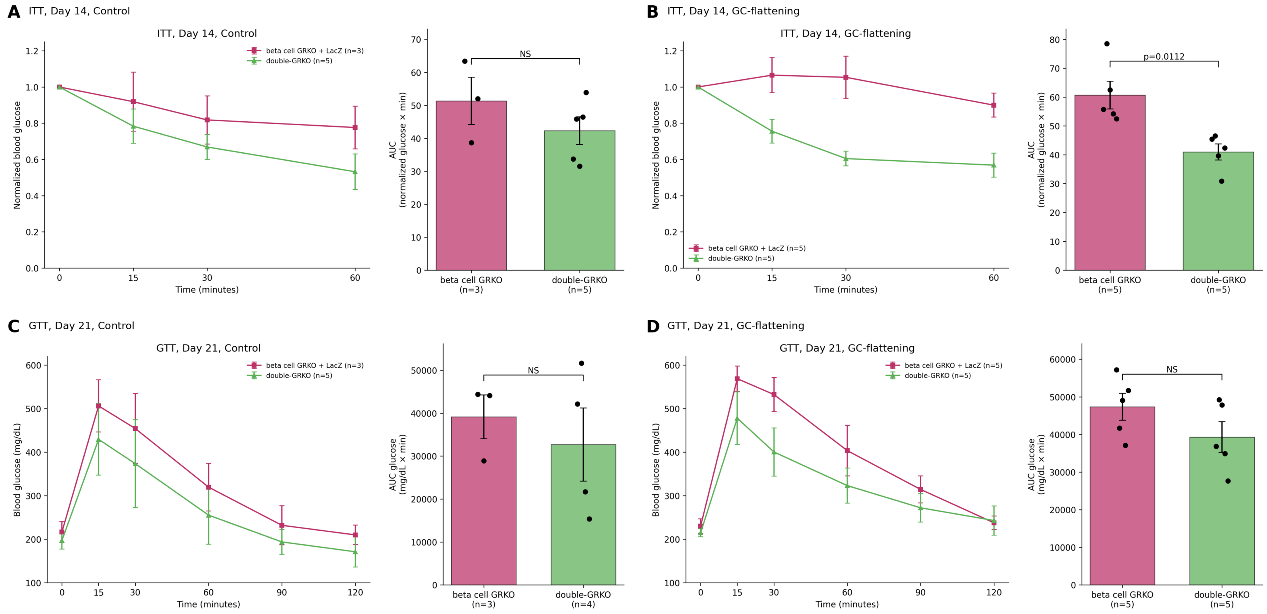


**ESM Fig. 5. AAV8-TBG-LacZ controls support the metabolic effects of hepatocyte GR deletion in double-GRKO mice.**

(A, B) Insulin tolerance tests (ITTs) at Day 14 in beta cell GRKO mice that received AAV8-TBG-LacZ (LacZ) or AAV8-TBG-iCre (double-GRKO), under control (A) and GC-flattening (B) conditions. Blood glucose is expressed relative to baseline, and area-under-the-curve (AUC) values are shown at right. (C, D) Glucose tolerance tests (GTTs) at Day 21 in beta cell GRKO mice that received AAV8-TBG-LacZ or AAV8-TBG-iCre, under control (C) and GC-flattening (D) conditions. AUC values are shown at right. Blood glucose values that exceeded the glucometer detection limit were assigned a value of 600 mg/dl for plotting and AUC calculation. Under control conditions, n = 3 for LacZ and n = 5 for double-GRKO mice in the ITT, and n = 3 for LacZ and n = 4 for double-GRKO mice in the GTT AUC analysis. Under GC-flattening conditions, n = 5 for LacZ and n = 5 for double-GRKO mice. AUC measurements were analysed by two-way ANOVA with treatment and AAV/genotype as factors followed by Šídák-adjusted planned comparisons of AAV8-TBG-LacZ versus AAV8-TBG-iCre within each treatment condition. Where pairwise statistical comparisons are indicated on the panels, only statistically significant comparisons are shown. Nonsignificant comparisons are not displayed. Panel B shows the exact Šídák-adjusted p value (p=0.0112). These controls support that the increased insulin sensitivity observed in double-GRKO mice during GC-flattening is associated with hepatocyte GR deletion rather than with nonspecific effects of AAV administration.
